# Multi-scale semi-supervised clustering of brain images: deriving disease subtypes

**DOI:** 10.1101/2021.04.19.440501

**Authors:** Junhao Wen, Erdem Varol, Aristeidis Sotiras, Zhijian Yang, Ganesh B. Chand, Guray Erus, Haochang Shou, Ahmed Abdulkadir, Gyujoon Hwang, Dominic B. Dwyer, Alessandro Pigoni, Paola Dazzan, Rene S. Kahn, Hugo G. Schnack, Marcus V. Zanetti, Eva Meisenzahl, Geraldo F. Busatto, Benedicto Crespo-Facorro, Romero-Garcia Rafael, Christos Pantelis, Stephen J. Wood, Chuanjun Zhuo, Russell T. Shinohara, Yong Fan, Ruben C. Gur, Raquel E. Gur, Theodore D. Satterthwaite, Nikolaos Koutsouleris, Daniel H. Wolf, for the Alzheimer’s Disease Neuroimaging Initiative, Christos Davatzikos

**Author notes:** Corresponding authors: Junhao Wen, PhD –, Christos Davatzikos, PhD – Christos. 3700 Hamilton Walk, Philadelphia, PA 19104. Data used in preparation of this article were obtained from the Alzheimer’s Disease Neuroimaging Initiative (ADNI) database (adni.loni.usc.edu). As such, the investigators within the ADNI contributed to the design and implementation of ADNI and/or provided data but did not participate in analysis or writing of this report. A complete listing of ADNI investigators can be found at: http://adni.loni.usc.edu/wp-content/uploads/how_to_apply/ADNI_Acknowledgement_List.pdf.

## Abstract

Disease heterogeneity is a significant obstacle to understanding pathological processes and delivering precision diagnostics and treatment. Clustering methods have gained popularity for stratifying patients into subpopulations (i.e., subtypes) of brain diseases using imaging data. However, unsupervised clustering approaches are often confounded by anatomical and functional variations not related to a disease or pathology of interest. Semi-supervised clustering techniques have been proposed to overcome this and, therefore, capture disease-specific patterns more effectively. An additional limitation of both unsupervised and semi-supervised conventional machine learning methods is that they typically model, learn and infer from data using a basis of feature sets pre-defined at a fixed anatomical or functional scale (e.g., atlas-based regions of interest). Herein we propose a novel method, “Multi-scAle heteroGeneity analysIs and Clustering” (MAGIC), to depict the multi-scale presentation of disease heterogeneity, which builds on a previously proposed semi-supervised clustering method, HYDRA. It derives multi-scale and clinically interpretable feature representations and exploits a double-cyclic optimization procedure to effectively drive identification of inter-scale-consistent disease subtypes. More importantly, to understand the conditions under which the clustering model can estimate true heterogeneity related to diseases, we conducted extensive and systematic semi-simulated experiments to evaluate the proposed method on a sizeable healthy control sample from the UK Biobank (N=4403). We then applied MAGIC to imaging data from Alzheimer’s disease (ADNI, *N*=1728) and schizophrenia (PHENOM, *N*=1166) patients to demonstrate its potential and challenges in dissecting the neuroanatomical heterogeneity of common brain diseases. Taken together, we aim to provide guidance regarding when such analyses can succeed or should be taken with caution. The code of the proposed method is publicly available at https://github.com/anbai106/MAGIC.

**Highlights:**

- We propose a novel multi-scale semi-supervised clustering method, termed MAGIC, to disentangle the heterogeneity of brain diseases.
- We perform extensive semi-simulated experiments on large control samples (UK Biobank, *N*=4403) to precisely quantify performance under various conditions, including varying degrees of brain atrophy, different levels of heterogeneity, overlapping disease subtypes, class imbalance, and varying sample sizes.
- We apply MAGIC to MCI and Alzheimer’s disease (ADNI, *N*=1728) and schizophrenia (PHENOM, *N*=1166) patients to dissect their neuroanatomical heterogeneity, providing guidance regarding the use of the semi-simulated experiments to validate the subtypes found in actual clinical applications.

**Graphical abstract:** 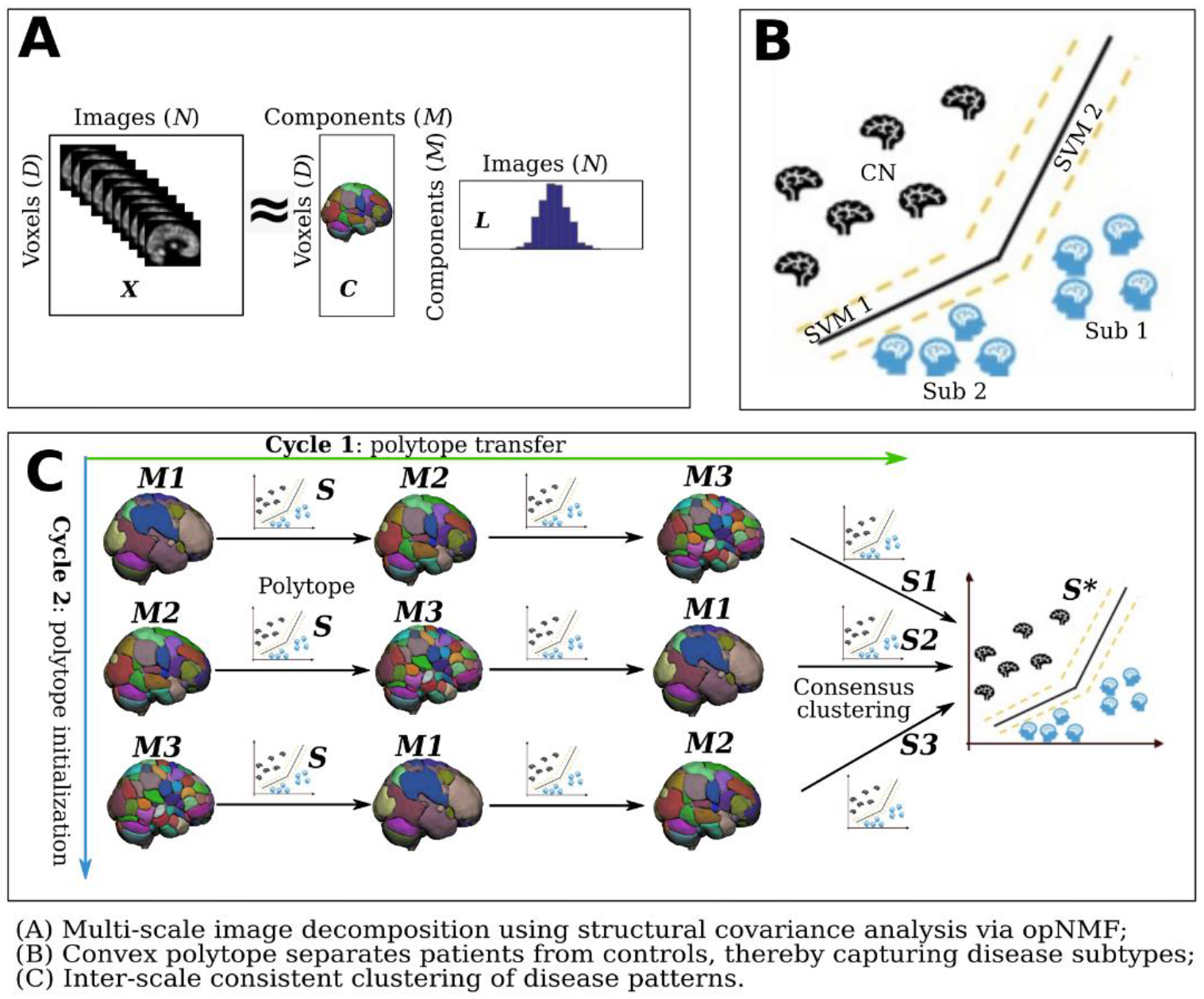

## 1. Introduction

Statistical and machine learning (ML) methods have been widely applied to neuroimaging data to derive disease-specific imaging signatures (Davatzikos, 2019). Voxel-based analysis (VBA) techniques generally involve performing independent mass univariate statistical tests on all voxels (Ashburner et al., 1998; Ashburner and Friston, 2000; Davatzikos et al., 2001; Friston et al., 1994), aiming to unveil detailed spatial maps of brain structures that are associated with clinical variables of interest. However, VBA approaches suffer from limited statistical power since they ignore multivariate data interactions. In contrast, multivariate pattern analysis (MVPA) techniques have gained traction due to their ability to capture complex multivariate interactions in data. Classical multivariate models, such as support vector machine (SVM), have been extensively utilized in the neuroimaging community to reveal imaging signatures for several brain diseases and disorders (Cuingnet et al., 2011; Ecker et al., 2010; Gaonkar and Davatzikos, 2013; Habes et al., 2016; Koutsouleris et al., 2015; Lao et al., 2004; Rathore et al., 2017; Samper-González et al., 2018; Varol et al., 2018). More recently, highly nonlinear and multivariate deep learning models have also been applied to brain modeling (Bashyam et al., 2020; Schulz et al., 2020b; Wen et al., 2020a). However, due to possible over-fitting, these models’ interpretability and generalizability in low sample size regimes have been under scrutiny.

Whether performing mass univariate or multivariate analysis, it is typically assumed that a relatively pure pathological pattern exists in the disease population. The disease signature is often presented via a voxel-wise or region of interest (ROI)-wise statistical map of the case-control group differences, i.e., between healthy controls (CN) and patients (PT). However, in nature, disease effects are commonly heterogeneously presented across different subpopulations due to the diversity of underlying risk factors. Such model assumption violations may cause the statistical learning to yield underpowered or false-positive results (Dwyer et al., 2018). Tackling this issue is of great importance given ample evidence of disease heterogeneity (Murray et al., 2011; Noh et al., 2014; Whitwell et al., 2007) and increasing appreciation that this may undermine the precision of clinical treatment guidelines and obscure research findings (Insel and Cuthbert, 2015).

Disentangling disease heterogeneity elucidates the underlying pathological mechanisms and potentially enables clinicians to offer targeted treatment options to different patient subpopulations. Nonlinear methods, such as deep neural networks, implicitly handle heterogeneity. However, there still exists a gap between these models and human interpretability, especially for clinicians who frequently seek discrete disease subtypes (Miotto et al., 2018). Thus, many recent efforts to discover the heterogeneous nature of brain diseases have investigated different clustering algorithms (Chand et al., 2020; Dong et al., 2016a, 2016b; Dwyer et al., 2018; Ezzati et al., 2020; Filipovych et al., 2012; Honnorat et al., 2019; Jeon et al., 2019; Jung et al., 2016; Lubeiro et al., 2016; Nettiksimmons et al., 2014; Ota et al., 2016; Pan et al., 2020; Park et al., 2017; Planchuelo-Gómez et al., 2020; Poulakis et al., 2020, 2020, 2018; Sugihara et al., 2016; Ten Kate et al., 2018; Varol et al., 2017; Young et al., 2018; Zhang et al., 2016). These methods can be divided into two categories depending on whether the clustering algorithm is unsupervised or semi-supervised^b^. Unsupervised clustering techniques, such as K-means (Hartigan and Wong, 1979), hierarchical clustering (Day and Edelsbrunner, 1984), and non-negative matrix factorization (NMF) (Lee and Seung, 2001), aim to directly cluster the patients based on their demographic information, clinical presentation, or imaging biomarkers. However, the results of these methods have often been confounded by non-pathologic processes, such as demographics. To cope with these covariate confounds, semi-supervised clustering methods (Dong et al., 2016a; Varol et al., 2017) leverage the group-level information and attempt to nullify the effect of nuisance variables. These methods generate clusters based on the pattern differences between the CN population and the subpopulations of patients (i.e., subtypes/clusters), hypothesizing that each pattern represents a distinct disease dimension or subtype. The main limitation of this family of methods is that they usually seek subtypes on a single scale set of features (e.g., atlas-based ROIs, voxels, networks), which makes the result heavily dependent on the level of granularity of the feature space. However, there has been abundant evidence that the brain is fundamentally constructed by multi-scale entities (Bassett and Siebenhühner, 2013; Betzel and Bassett, 2017). Therefore, it is beneficial to analyze disease heterogeneity on multiple spatial scales and seek a compatible clustering solution across scales, which will potentially better align with the brain’s multi-scale nature.

Despite the fact that these clustering analyses have always led to a cluster solution, there are no clear guidelines enabling the validity of the cluster solution to be determined, presumably due to the lack of the ground truth in clustering problems or the “curse of dimensionality” in brain imaging settings. A previous study (Varol et al., 2017) designed simulation experiments to validate the proposed model. However, the simulation data were generated by adding noise in the low-dimensional feature space under a specific distribution (i.e., Gaussian distribution), which was far less realistic than actual neuroimaging data. Thus, a more sophisticated and systematic simulation is needed to understand the conditions under which clustering succeeds or fails with high-dimensional brain imaging data. Specifically, in the current work, we performed an extensive and systematic evaluation of clustering performance using a large healthy control sample (UK Biobank, *N*=4403) in a semi-simulated setting. The term semi-simulated here refers to the fact that brain heterogeneity may stem from various sources, and the simulation was performed with data from real healthy control individuals. We simulated the heterogeneity due to disease effects by imposing abnormalities (i.e., increasing or decreasing voxel intensity) on specific regions of tissue images. Notably, the heterogeneity caused by normative brain aging was inevitably retained in the original data because this is biologically realistic and contributs to the semi-simulated variability (refer to Section 4.2 for more details). With known ground truth for the number of clusters (*k*) and the cluster/subtype membership assignment, we quantitatively investigated the clustering model’s performance under a variety of conditions, including varying degrees of brain atrophy, different levels of heterogeneity, overlapping disease subtypes, class imbalance, and varying sample sizes.

This work is a comprehensive extension of our preliminary results presented in Medical Image Computing and Computer Assisted Interventions (MICCAI) 2020 (Wen et al., 2020b). The contribution is two-fold. First, to address the aforementioned multi-scale limitations, we propose a data-driven and multi-scale semi-supervised method termed MAGIC for “Multi-scAle heteroGeneity analysIs and Clustering”. Specifically, MAGIC extracts multi-scale features, from coarse to fine granularity, via orthogonal projective non-negative matrix factorization (opNMF) applied for varying scales (i.e., number of components). opNFM has been a very effective unbiased, data-driven method for extracting biologically interpretable and reproducible feature representations in the context of neuroimaging datasets (Sotiras et al., 2015), leading to disease subtypes in an explainable space (Schulz et al., 2020a). A convex polytope classifier, based on principles of the method in (Varol et al., 2017), is applied to these multi-scale features through a double-cyclic optimization procedure to yield robust clusters that are consistent across different scales. Secondly, the results of our semi-simulated experiments allow us to compare MAGIC with previous standard clustering methods and provide future clustering analysis guidelines. Specifically, applying the proposed method to Alzheimer’s disease (AD) and mild cognitive impairment (MCI) and schizophrenia (SCZ) patients provides greater confidence regarding the validity of the subtypes claimed in actual clinical applications.

We organize the remainder of the paper as follows. In Section 2, we provide the details of the proposed algorithm. Section 3 details the primary datasets and image preprocessing steps. Section 4 presents the results of the experiments. Section 5 concludes the paper by discussing our main observations, method limitations, and future directions.

## 2. Methods

MAGIC builds upon the HYDRA formulation (Varol et al., 2017) and opNMF algorithms (Sotiras et al., 2015) to yield an inter-scale-consistent clustering solution. It generates an interpretable and spatially adaptive multi-scale representation via opNMF, which drives semi-supervised clustering. The schematic diagram of MAGIC is shown in Fig. 1.

**Figure 1.**
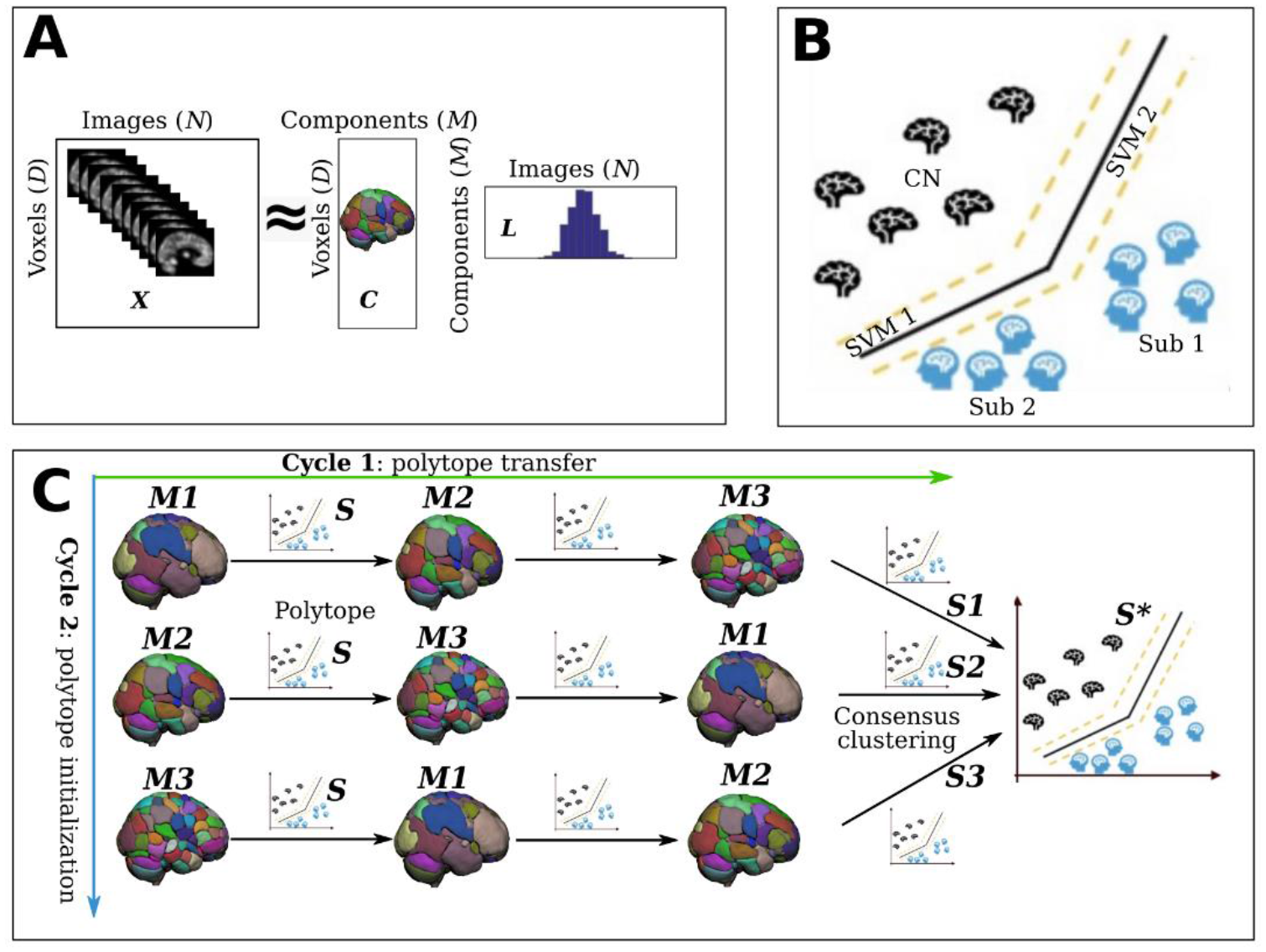
Schematic diagram of the MAGIC algorithm. MAGIC first generates multi-scale feature representations of the brain anatomy from coarse to fine resolutions and then cyclically solves semi-supervised clustering subproblems with each of these feature representations. Generally, it consists of three key components. A) opNMF enables the extraction of multi-scale, biologically interpretable feature representations in a data-driven manner. B) max-margin multiple SVM classifiers are utilized to construct a nonlinear polytope for simultaneous classification and clustering. In this fashion, the patients’ subtypes or subpopulations are clustered based on their distance from the polytope. C) the double-cyclic optimization procedure is adopted to fuse the knowledge from multi-scale features for inter-scale consistent clustering solutions. Specifically, the cluster polytope is first initialized at a specific representation scale. After optimization, the cluster polytope is transferred to the next representation scale, allowing the clustering routine to be guided by all anatomical scales. Furthermore, the polytope initialization is performed at different anatomical scales to further remove bias from the clustering solutions. Lastly, the resulting multi-scale clustering solutions are fused through consensus clustering to yield a final stable subtype membership assignment. ***X***: input matrix; ***C***: component matrix; ***L***: loading coefficient matrix; CN: healthy control; Sub: subtype; ***M***: number of components. ***S*** is the initial polytope solution. ***S1***, ***S2***, and ***S3*** are the fine-tuned polytope for different initialization models, and ***S\**** is the final polytope after the consensus clustering procedure.

We detail the mathematical formulation of the optimization routine in the following subsections. To establish notation, let *N* denote the number of subjects and *D* the number of voxels in each image. We denote the data as a matrix ***X*** that is organized by arranging each image as a vector per column (***X*** = [***x****_1_*, …, ***x****_N_*], ***x****_i_* ∈ *R^D^*). We use binary labels to distinguish the patient and control groups, where 1 represents patients (PT) and - 1 means healthy controls (CN) (i.e., ***y*** ∈ {−1, 1}^*N*^). For subtype results, the subtype membership matrix (a.k.a., polytope) is denoted as ***S***∈ *R^N^* ^x^ *^k^* before consensus clustering and ***S\**** as the final subtype membership matrix after consensus clustering.

### 2.1. Multi-scale feature extraction via orthogonal projective non-negative matrix factorization

MAGIC utilizes opNMF, an unsupervised representation learning algorithm, to extract multi-scale and interpretable anatomical components covering the whole brain. The number of components (*M*) is optimized in opNMF and controls the granularity of the anatomical components (e.g., opNMF components at different granularities can be seen in Fig. 1C).

The opNMF aims to represent the input matrix ***X*** as a rank-*M* matrix that is the product of two non-negative matrices: i) ***C***, termed as the component matrix, captures the groups of voxels that covary most and offers an interpretable anatomical parcellation (***C*** = [***c****_1_*, …, ***c****_M_*], ***c****_i_* ∈*R^D^*), and ii) ***L*** ∈*R^M^*^x^*^N^*, termed as the loading coefficient matrix, captures the amount of each spatial component that makes up each subject. The opNMF objective is to be minimized as follows:

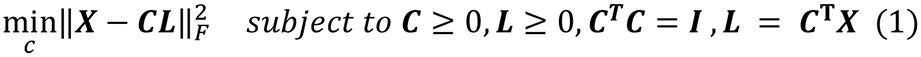

This formulation differs from the standard NMF in that the loading coefficient matrix is obtained by projecting the input data ***X*** to the estimated component matrix ***C*** (i.e., ***L*** = ***C*^T^*X***), and the orthogonality constraint is imposed on the component matrix (***C*^T^*C*** = ***I***, where ***I*** denotes the identity matrix). Therefore, the opNMF searches only the parameters of the component matrix during optimization (Zhirong Yang and Oja, 2010).

The solution of minimizing the abovementioned objective is a non-convex problem and can be achieved by iteratively updating the multiplicative rule proposed in (Zhirong Yang and Oja, 2010):

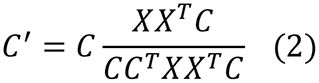

Please refer to (Sotiras et al., 2015) for more details about opNMF and (Zhirong Yang and Oja, 2010) for convergence analyses. Once the algorithm converges, we recover the loading coefficients by the projective step: ***L*** = ***C***^**T**^***X***. Moreover, this property allows us to readily apply the trained model to external unseen data.

### 2.2. Max-margin multiple SVM classifiers for clustering

Once the high dimensional imaging data is reduced to a lower-dimensional representation using opNMF, we apply the HYDRA algorithm (Varol et al., 2017) on the set of loading coefficients, ***L*** ∈ *R^M^*^x^*^N^* and the corresponding set of diagnostic labels ***y*** ∈ {−1, 1}^*N*^ to perform clustering of the patients.

The HYDRA algorithm utilizes multiple large margin classifiers (e.g.., *k* SVMs) to estimate a nonlinear polytope that separates the two classes with maximized distance (or margin) from the decision boundaries for each sample, thus simultaneously serving for classification and clustering. The fundamentals of the HYDRA algorithm are presented in supplementary eMethod 1. Please refer to (Varol et al., 2017) for more details. In general, this algorithm solves for a convex polytope classification boundary that discriminates patients from controls with a maximum margin. In essence, the polytope is composed of the *k* hyperplanes of the *k* linear SVMs, and each face corresponds to one subtype/cluster. The objective of maximizing the polytope’s margin can be summarized as:

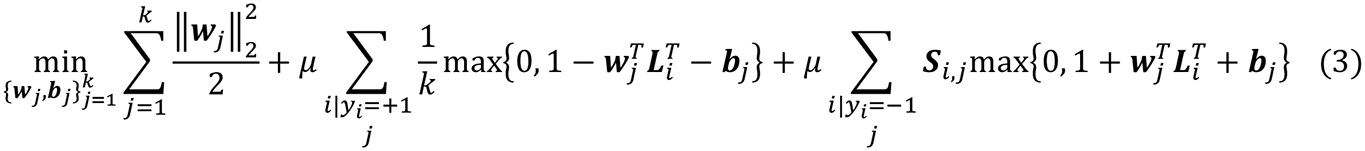

where ***w***_*j*_ and ***b***_*j*_ are the weight and bias for each hyperplane, respectively. *μ* is a penalty parameter on the training error, and ***S*** *is* the subtype membership matrix of dimension *NxK* containing information regarding whether a sample *i* belongs to subtype *j*. In general, this optimization problem is non-convex and is jointly optimized by iterating on solving for the polytope faces’ parameters using standard SVM solvers (Chang and Lin, 2011) and solving for the cluster memberships as follows:

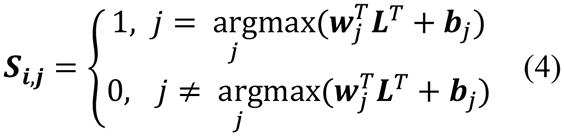

### 2.3. Double-cyclic optimization procedure for scale-independent subtypes

MAGIC optimizes the clustering objective, i.e., Eq. 3, for each anatomical scale as a sub-optimization problem. To fuse the multi-scale clustering solutions and enforce the clusters to be scale-independent, MAGIC adopts a double-cyclic procedure that transfers and fine-tunes the subtype membership matrix (***S***) between different scales of features, i.e., solving the sub-optimization problems with the single-scale feature representation in a loop (Fig. 1).

The double-cyclic fine-tuning procedure aims to offer scale-independent clustering solutions across multi-scale features. Cycle 1 (Fig. 1C components *M*1, *M*2, and *M*3 in a row) aims to derive a clustering solution that is informed by features across all scales. This is achieved by iteratively solving Eq. 2 using features derived at different scales. Specifically, the clustering membership matrix ***S*** is first solved for a particular set of features. It is then transferred to the next block, where it is used as initialization for fine-tuning driven by features from a different scale. This procedure is repeated till features from all anatomical scales have been used to inform the final clustering membership matrix (***S1*** in Fig. 1C). Since each optimization cycle starts at a pre-determined anatomical scale, an additional Cycle 2 (Fig. 1C components *M*1, *M*2, and *M*3 in a column) is executed using all different anatomical scales to initialize the model. This eliminates any initialization biases (***S1***, ***S2***, and ***S3*** in Fig. 1C) and results in multiple clustering solutions. To determine the final subtype assignment (***S\**** in Fig. 1C), we perform consensus clustering. Consensus is achieved by grouping together samples that are assigned to the same cluster across the solutions estimated as part of Cycle 2 (Varol et al., 2017). Precisely, we first compute a co-occurrence matrix based on the clustering results of Cycle 2 and then use it to perform spectral clustering (Ng et al., 2001).

MAGIC can be directly applied to unseen external data with the following procedure. First, opNMF is not required to be retrained to unseen data because multi-scale feature extraction can be achieved via the projection ***L*** = ***C***^**T**^***X***. Subsequently, each single-scale feature is fit to each polytope (***S1***, ***S2***, and ***S3*** in Fig. 1) to derive the single-scale clustering solution. Finally, a similar consensus procedure is used to derive the final membership (***S\**** in Fig. 1).

## 3. Materials

### 3.1. Datasets

Three datasets are used in the current study: the UK Biobank (UKBB) study (Miller et al., 2016), the Alzheimer’s Disease Neuroimaging Initiative (ADNI) study (Petersen et al., 2010), and the Psychosis Heterogeneity Evaluated via Dimensional Neuroimaging (PHENOM) study (Chand et al., 2020; Rozycki et al., 2018; Satterthwaite et al., 2010; Schnack et al., 2014, 2014; Wolf et al., 2014; Wood et al., 2001; T. Zhang et al., 2015; Zhu et al., 2016; Zhuo et al., 2016).

The UKBB is a dataset of approximately 500,000 UK adults sampled via population-based registries (http://www.ukbiobank.ac.uk). Participants were recruited from across the United Kingdom, and initial enrolment was carried out from 2006 to 2010. Participants provided socio-demographic, cognitive, and medical data via questionnaires and physical assessments. Starting in 2014, a subset of the original sample later underwent brain magnetic resonance imaging (MRI). The UKBB data used in this work comprises 4403 CN participants whose T1-weighted (T1w) MRI was collected using Siemens 3T Skyra. The parameters of the 3D MPRAGE sequences are as follows: resolution=1.0×1.0×1.0 mm; field-of-view=256 mm x256 mm; TR = 2000 ms; TE = 2.01 ms; TI = 880 ms; slices = 208; flip angle = 8 degrees (Miller et al., 2016).

The ADNI was launched in 2003 as a public-private partnership (https://www.adni-info.org/). The primary goal of ADNI has been to test whether serial MRI, positron emission tomography (PET), other biological markers, and clinical and neuropsychological assessment can be combined to measure the progression of MCI and early AD. The ADNI dataset used in our experiments comprises 1728 participants from ADNI 1, 2, 3, and GO, for whom a T1w MRI was available at baseline: 339 AD, 541 CN, and 848 MCI were finally included. ADNI T1w images were performed both on 1.5T and 3T scanners with similar protocol parameters: 256×256 matrix; voxel size=1.2×1.0×1.0 mm; TI=400 ms; TR=6.98 ms; TE=2.85 ms; flip angle=11°.

The PHENOM dataset is an international consortium spanning five continents to better understand neurobiological heterogeneity in schizophrenia. The consortium aims to delineate schizophrenia brain subtypes with large sample sizes, enriched sample heterogeneity, and methodological advances that generalize across disparate sites and ethnicities. The PHENOM dataset used in this study includes 1166 participants (583 CN, and 583 SCZ patients). In the current study, we included T1w images from eight sites of the PHENOM consortium with diverse imaging protocols.

These datasets are described in detail in supplementary eMethod 2. Table 1 summarizes the basic demographics of all participants from the three datasets.

**Table 1.**
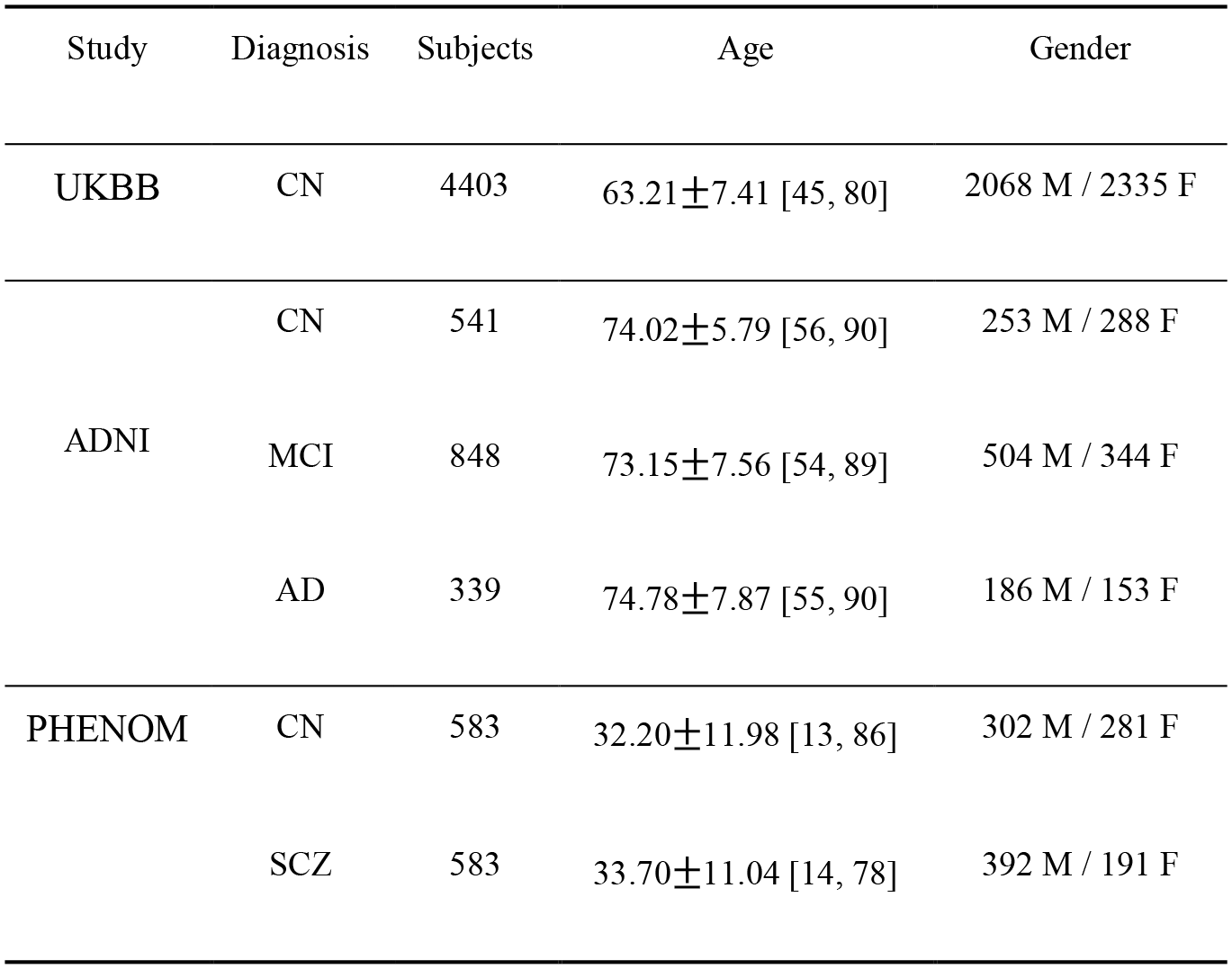
Summary of participant demographics for UKBB, ADNI, and PHENOM datasets. Values for age are presented as mean ± SD [range]. M: male, F: female.

### 3.2. Image preprocessing

Raw T1w MRIs were quality checked for motion, image artifacts, or restricted field-of-view. Images passing this quality check (QC) were corrected for magnetic field inhomogeneity (Tustison et al., 2010). A robust multi-atlas label fusion-based method, MUSE (Doshi et al., 2016), was applied for tissue segmentation of the brain. Voxel-wise regional volumetric maps (RAVENS) (Davatzikos et al., 2001) were generated for grey matter (GM) tissues by registering skull-stripped images to a population-based template residing in the MNI-space using a deformable registration method (Ou et al., 2011). Another QC procedure was performed to control the quality of the images further. Specifically, the images were first checked by manually evaluating for pipeline failures (e.g., poor brain extraction, tissue segmentation, and registration errors). Furthermore, a second-step automated procedure automatically flagged images based on outlying values of quantified metrics (i.e., ROI values), and those flagged images were re-evaluated.

## 4. Experiments and results

We first validated the proposed model using semi-simulated data in which we knew the ground truth for the number of clusters (*k*) and subtype membership assignment. In this setting, we quantitatively assessed how several key components influenced the clustering performance and compared our method’s performance to other common clustering approaches. Finally, we applied MAGIC to real clinical datasets for dissecting the heterogeneity of AD plus MCI and SCZ. In our experiments, different nuisance variables (i.e., age and sex or site) were controlled with a linear regression model in MAGIC for the multi-scale features. Specifically, the *beta* was estimated on healthy control subjects and then applied to all populations.

### 4.1. Evaluation strategy

We adopted a cross-validation (CV) procedure with repeated and stratified random splits for 100 repetitions to determine the appropriate number of clusters. Specifically, during each repetition, 80% of the data was for training. The “optimal” number of clusters was guided by the clustering stability across the 100 repetitions. The Adjusted Rand index (ARI) was used for that purpose, which we denoted as ARIs during CV (ARI_CV). Moreover, for simulation experiments, where the ground truth for subtype membership was known, ARI was also used to quantify the clustering performance, referred to as ARIs for ground truth (ARI_GT).

After obtaining the assignment of subtype membership, we performed voxel-wise group comparisons for RAVENS GM maps between each subtype with CNs using the *3dttest++* program (Cox et al., 2017) in AFNI (Cox, 1996) to detect the distinct neuroanatomical patterns of the corresponding subtypes. The two-sample t-test T-value map of AFNI was further converted to a P-value map applying correction for multiple comparisons with the Benjamini-Hochberg procedure. Effect sizes can for some purposes be more useful than P-values, since P-values are highly dependent on the sample size (Sullivan and Feinn, 2012). Thus, we calculated the effect size, Cohen’s *f*2 (Selya et al., 2012), for voxels that are significantly different between subtypes after adjusting the confounding covariates (i.e., age and sex). We chose Cohen’s *f*2 over Cohen’s *d* because the formulation of Cohen’s *f*2 takes into account the confounding covariates in a general linear model set-up, whereas Cohen’s *d* is simply the mean difference of two groups divided by the pooled standard deviation. We present the voxel-wise effect size maps to delineate the subtypes’ neuroanatomical patterns for all experiments. For reference, Cohen’s *f*2 ≥ 0.02, ≥ 0.15, and ≥ 0.35 represent small, medium, and large effect sizes, respectively (Selya et al., 2012).

### 4.2. Experiments using UKBB semi-simulated data

The UKBB RAVENS GM maps were used to generate semi-simulated data. We first divided all CN subjects (*N*=4403) into pre-defined number of splits. Part of the splits was regarded as the true CN, and the remainder (i.e., pseudo-PT) was further divided into another number of splits for subtype simulations. The sample size of each subtype was balanced. Brain atrophy was then imposed onto RAVENS maps of each of the subtypes within different patterns. To simplify the simulation, we assume that patterns across the *k* subtypes are orthogonal with each other (we further tested the influence of overlapping patterns between subtypes). These regions were chosen *a priori* based on the segmentation image of the template image in the MNI space. Different choices for the number of subtypes (*k*) and atrophy strength level (ASL) were tested. For instance, for experiments with *k*=2 and ASL=0.1, voxel intensity values inside the two pre-defined patterns were reduced by 10% compared to their original values. Moreover, the ASL varied by ±2% across images to add randomness. In total, nine experiments were performed and summarized in Table 2. The ground truth of the pre-defined atrophy patterns of each subtype is shown in Fig. 3 (i.e., the first column). During simulation, we ensured that the subtype groups did not significantly differ in sex and age. Of note, the UKBB subjects were primarily diagnosed as neurodegenerative-speaking healthy controls, but they were also self-reported for various comorbidities (see https://biobank.ndph.ox.ac.uk/ukb/field.cgi?id=41202). Therefore, age or comorbidity-related heterogeneity already exist in the original data. This setting is more realistic because heterogeneity caused by brain aging and pathologies often intertwine with each other.

**Table 2.** Summary of the original semi-simulated experiments. The number of subjects for each group is shown in parentheses. ASL: atrophy strength level; *k*: the number of clusters. Sub: Subtype.

| Experiment | Subtype and sample size |
| --- | --- |
| $k=2$ & ASL=0.1 | CN (2201), Sub1 (1101), Sub2 (1101) |
| $k=2$ & ASL=0.2 | CN (2201), Sub1 (1101), Sub2 (1101) |
| $k=2$ & ASL=0.3 | CN (2201), Sub1 (1101), Sub2 (1101) |
| $k=3$ & ASL=0.1 | CN (1103), Sub1 (1100), Sub2 (1100), Sub3 (1100) |
| $k=3$ & ASL=0.2 | CN (1103), Sub1 (1100), Sub2 (1100), Sub3 (1100) |
| $k=3$ & ASL=0.3 | CN (1103), Sub1 (1100), Sub2 (1100), Sub3 (1100) |
| $k=4$ & ASL=0.1 | CN (883), Sub1 (880), Sub2 (880), Sub3 (880), Sub4 (880) |
| $k=4$ & ASL=0.2 | CN (883), Sub1 (880), Sub2 (880), Sub3 (880), Sub4 (880) |
| $k=4$ & ASL=0.3 | CN (883), Sub1 (880), Sub2 (880), Sub3 (880), Sub4 (880) |

In sum, we sought to compare MAGIC’s clustering performance to other unsupervised or semi-supervised clustering methods. Since the influence of confounds on clustering performance may vary under different conditions, our second goal was to test under what conditions MAGIC can discover i) the true membership of the subtypes, ii) the true number of clusters (*k*); iii) its simulated atrophy patterns, and iv) its severity of the abnormal patterns (i.e., voxel-wise effect size map).

#### 4.2.1. MAGIC discovers the correct number of clusters and corresponding simulated neuroanatomical patterns

MAGIC was able to discover the correct number of clusters for the following experiments: *k*=2 & ASL=0.1, 0.2 or 0.3 (Fig. 2A, B and C), *k*=3 & ASL=0.2 (Fig. 2E) or ASL=0.3 (Fig. 2F), and *k*=4 & ASL=0.3 (Fig. 2I). For other experiments, MAGIC failed to find the true *k* (Fig. 2D, G, and H), indicating that in the presence of high heterogeneity (K>2 or 3) and very subtle disease effect (10%-20%), the algorithm reaches a detection threshold.

**Figure 2.**
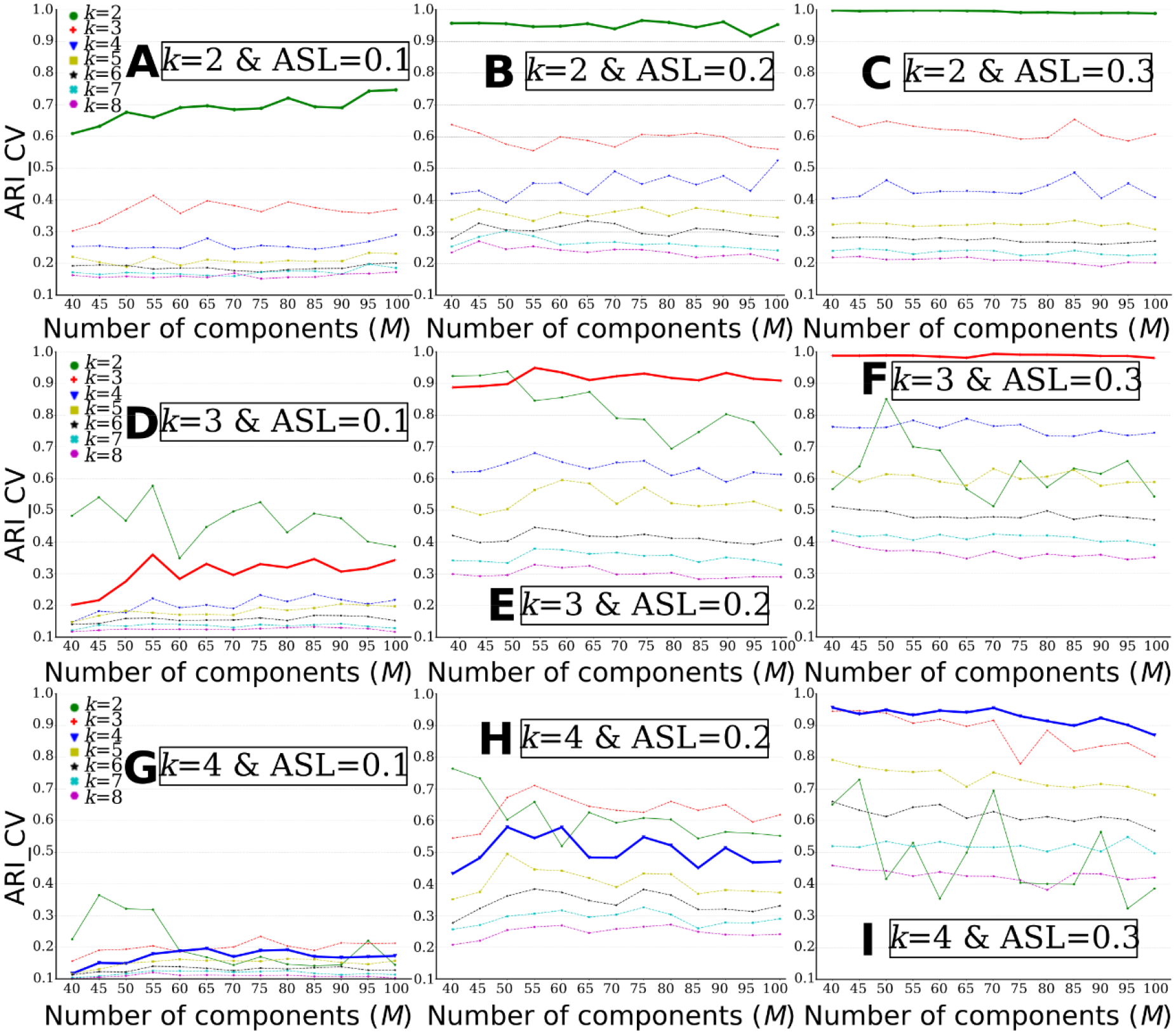
MAGIC finds the ground truth of the number of clusters (*k*) when the clustering conditions are favorable, i.e., higher ASL or lower *k*. The “optimal” *k* was determined by ARI_CV. A) *k*=2 & ASL=0.1; B) *k*=2 & ASL=0.2; C) *k*=2 & ASL=0.3; D) *k*=3 & ASL=0.1; E) *k*=3 & ASL=0.2; F) *k*=3 & ASL=0.3; G) *k*=4 & ASL=0.1; H) *k*=4 & ASL=0.2; I) *k*=4 & ASL=0.3. The bold lines represent the ground truth of *k* for each experiment.

Voxel-wise effect size maps were generated to demonstrate whether MAGIC can find the ground truth of neuroanatomical atrophy patterns of subtypes. Of note, the actual neuroanatomical patterns revealed by the voxel-wise maps include heterogeneity due to both the simulation effects (i.e., disease effect) and normative brain aging, e.g., those voxels without any simulation in the P-value mask map (Fig. 3). To further support the normative brain aging heterogeneity, we derived the voxel-wise effect size map for the original images (without any simulation) of subjects from Sub1 and healthy control groups in Fig. 3A (supplementary eFigure 1), which showed specific abnormality patterns with small effect sizes. Moreover, we quantitatively evaluated how well MAGIC can recover the simulated voxels. For that purpose, we proposed a simulation accuracy metric (ACC): the proportion of the number of voxels that passed the statistical significance in the P-value mask maps over the number of voxels in the ground truth mask that was masked by the population-based RAVENS GM tissue mask (Fig. 3).

**Figure 3.**
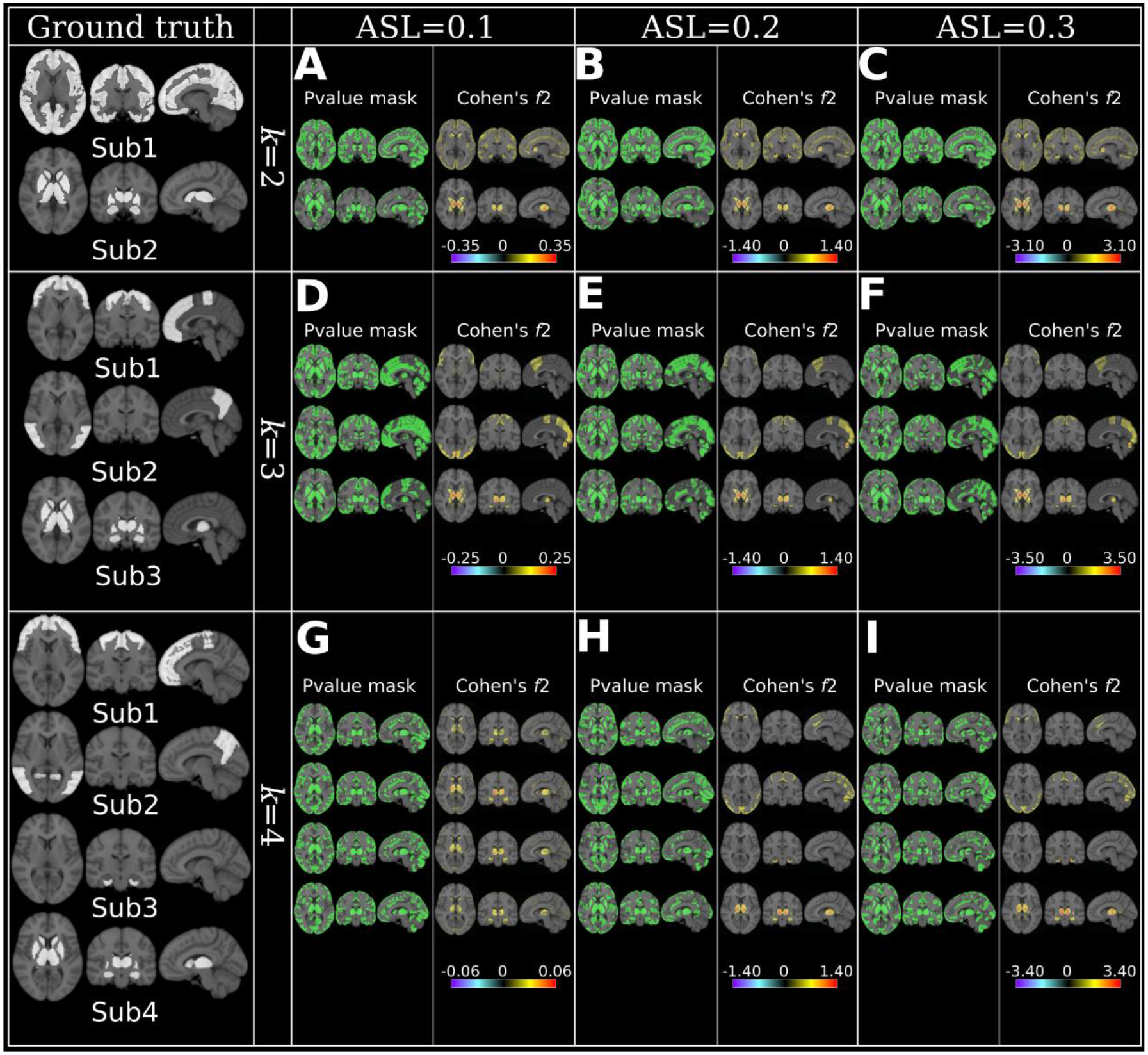
MAGIC finds the ground truth of subtype’s neuroanatomical patterns when the clustering conditions are favorable, i.e., higher ASL or lower *k*. Neuroanatomical patterns are displayed using effect size maps based on voxel-wise group comparisons between CN and subtypes. Positive values denote brain atrophy (CN > Sub), while negative values correspond to larger brain volume in subtypes (CN < Sub). The ground truth of the subtypes pattern is presented with a binary mask (white) for each *k* in the first column. Moreover, we proposed a simulation accuracy metric (ACC): the proportion of the number of voxels that passed the statistical significance in the P-value mask maps over the number of voxels in the ground truth mask that was masked by the population-based RAVENS GM tissue mask. A) *k*=2 & ASL=0.1, Sub1: ACC=0.77, Sub2: ACC=0.81; B) *k*=2 & ASL=0.2, Sub1: ACC=0.78, Sub2: ACC=0.84; C) *k*=2 & ASL=0.3, Sub1: ACC=0.78, Sub2: ACC=0.89; D) *k*=3 & ASL=0.1, Sub1: ACC=0.67, Sub2: ACC=0.70, Sub3: ACC=0.82; E) *k*=3 & ASL=0.2, Sub1: ACC=0.73, Sub2: ACC=0.74, Sub3: ACC=0.87; F) *k*=3 & ASL=0.3, Sub1: ACC=0.76, Sub2: ACC=0.72, Sub3: ACC=0.89; G) *k*=4 & ASL=0.1, Sub1: ACC=0.55, Sub2: ACC=0.54, Sub3: ACC=0.66, Sub4: ACC=0.59; H) *k*=4 & ASL=0.2, Sub1: ACC=0.66, Sub2: ACC=0.64, Sub3: ACC=0.65, Sub4: ACC=0.77; I) *k*=4 & ASL=0.3, Sub1: ACC=0.63, Sub2: ACC=0.66, Sub3: ACC=0.64, Sub4: ACC=0.79. For reference, Cohen’s *f*2 ≥ 0.02, ≥ 0.15, and ≥ 0.35 represent small, medium, and large effect sizes, respectively.

In short, MAGIC was able to find the ground truth for all experiments, except for *k*=4 & ASL=0.1 (Fig. 3G), in which small effects (Cohen’s *f*2<0.06) were detected in subcortical structures for all four subtypes. The voxels showing the largest effect sizes discovered by the effect size map were from the simulated regions. Furthermore, the P-value mask maps quantitatively showed that most of the simulated voxels could be detected by MAGIC (Fig. 3). Lastly, the effect size of the subtype patterns increased with increasing ASL (refer to the effect sizes in each row of Fig. 3).

#### 4.2.2. Comparison of MAGIC to other clustering methods

We compared MAGIC to other commonly used unsupervised clustering methods and HYDRA. Specifically, K-means is a vector quantification method that aims to partition the patient population into *k* clusters in which each participant belongs to the cluster with the nearest mean (Hartigan and Wong, 1979). GMM performs clustering by assuming that there are specific numbers of Gaussian distributions in patients, and each of these distributions belongs to one cluster (McLachlan and Basford, 1988). NMF aims to factorize the input matrix into two low-rank matrices with non-negative values. Intrinsically, the loading coefficient matrix conveys the clustering membership assignment (Lee and Seung, 2001). Lastly, the agglomerative hierarchical clustering (AHC) method is another unsupervised clustering method that seeks to build a hierarchy of clusters in a “bottom-up” fashion (Day and Edelsbrunner, 1984). Moreover, we fit the unsupervised methods and HYDRA with i) single-scale features (dotted curve lines in Fig. 4) and ii) multi-scale features (solid straight lines in Fig. 4) together for comprehensive comparisons, since MAGIC always take multi-scale features.

**Figure 4.**
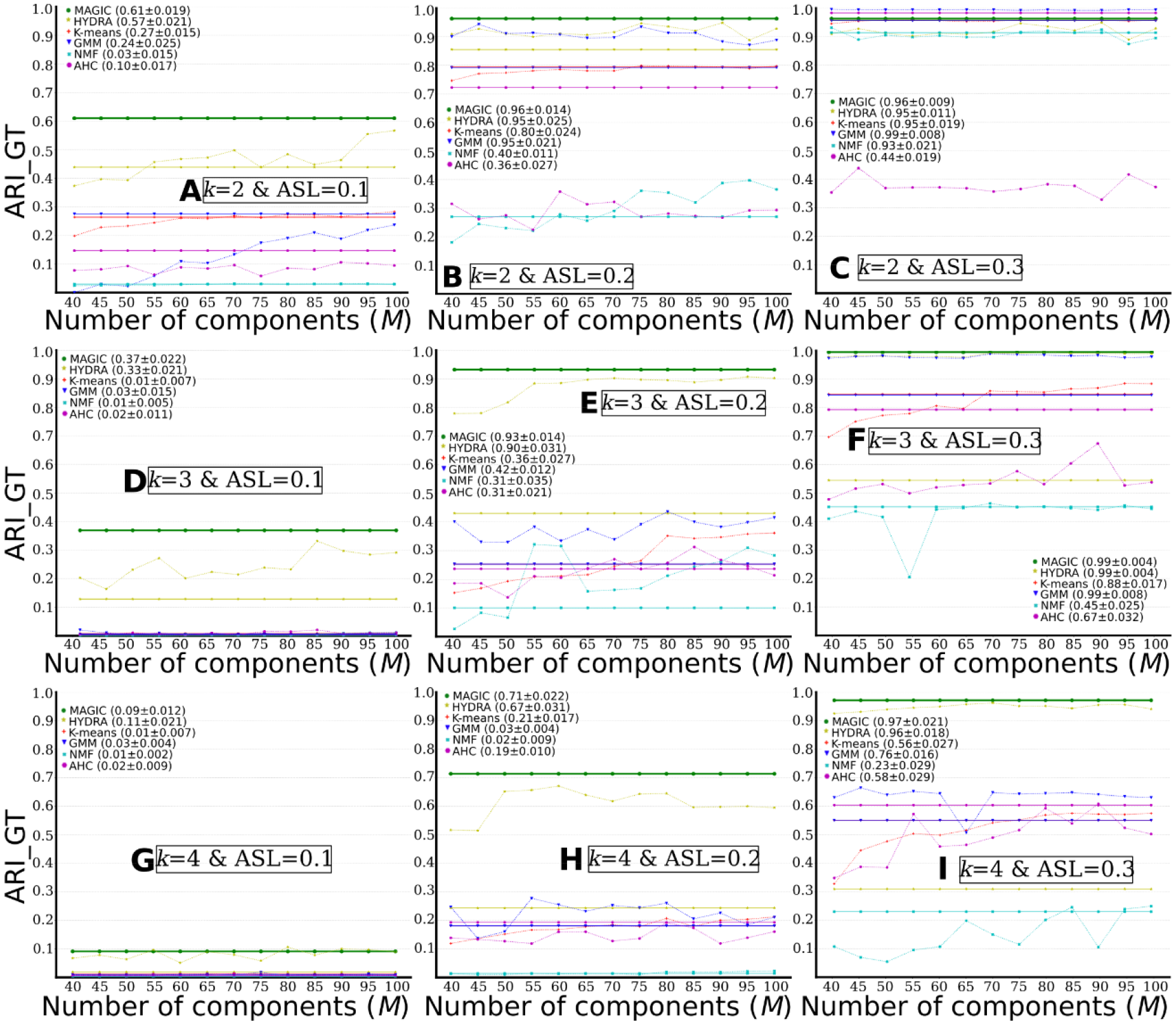
MAGIC outperforms other common clustering methods. Comparisons of clustering performance between different methods: MAGIC, HYDRA, K-means, GMM, NMF, and agglomerative hierarchical clustering (AHC) (*M*=40 to 100 with a step as 5). The solid straight lines show clustering results for models that take multi-scale features as input and are drawn over all *M*s only for visualization purposes. The dotted curve lines represent clustering results for models that take single-scale features as input. A) *k*=2 & ASL=0.1; B) *k*=2 & ASL=0.2; C) *k*=2 & ASL=0.3; D) *k*=3 & ASL=0.1; E) *k*=3 & ASL=0.2; F) *k*=3 & ASL=0.3; G) *k*=4 & ASL=0.1; H) *k*=4 & ASL=0.2; I) *k*=4 & ASL=0.3. We report the final consensus clustering performance (ARI_GT) for all models, together with the standard deviation of the 100-repetition clustering results during CV. For models using single-scale features, we show the results with the single-scale obtaining the highest ARI_GT.

As displayed in Fig. 4, MAGIC obtained slightly better clustering results than HYDRA and substantially outperformed all other unsupervised clustering methods (i.e., K-means, GMM, NMF, and agglomerative hierarchical clustering). Specifically, MAGIC obtained higher ARI_GTs for the following experiments: *k*=2 & ASL=0.1 (Fig. 4A), *k*=3 & ASL=0.1 or 0.2 or 0.3 (Fig. 4D, E and F), and *k*=4 & ASL=0.2 or 0.3 (Fig. 4H and I). All methods failed in clustering for experiment *k*=4 & ASL=0.1 (Fig. 4G). Furthermore, fitting all multi-scale features for HYDRA did not always perform better than the single-scale features and performed worse than MAGIC. Of note, fitting all multi-scales features (i.e., 910 features) for HYDRA took a much longer time to converge the model than single-scale HYDRA or MAGIC. For all experiments, we showed the consensus clustering performance and the standard deviation of the clustering performance across the 100 repetitions (Fig. 4). We decided not to report P-values because the “probability” of a false positive in this cross-validation scenario tends to be inflated. After all, no unbiased estimator of the correlation between the results obtained on the different repetitions exists (Nadeau and Bengio, 2003).

#### 4.2.3. Influence of the number of clusters

When the number of clusters *k* increased, MAGIC’s clustering performance gradually decreased (i.e., each column in Fig. 4 represents the three experiments with the same ASL), except for experiments *k*=4 & ASL=0.3. For ASL=0.1, the ARI_GTs are 0.610, 0.368 and 0.091 for *k*=2, 3 and 4, respectively. For ASL=0.2, the ARI_GT decreased from 0.960 to 0.934 and to 0.713 for *k*=2, 3 and 4, respectively. For ASL=0.3, the ARI_GTs are 0.994, 0.995 and 0.966 for *k*=2, 3 and 4, respectively.

#### 4.2.4. Influence of atrophy strength levels

With the increase of ASL, MAGIC’s clustering performance gradually improved (i.e., each row in Fig. 4 represents the three experiments with the same *k*). For *k*=2, the ARI_GTs are 0.610, 0.960 and 0.994 for ASL=0.1, 0.2 and 0.3, respectively. For *k*=3, the ARI_GT increased from 0.368 to 0.934 and to 0.995 for ASL=0.1, 0.2 and 0.3, respectively. For *k*=4, the ARI_GTs are 0.091, 0.713 and 0.966 for ASL=0.1, 0.2 and 0.3, respectively.

We visualized the subtypes/clusters in 2D space for all experiments using multidimensional scaling (Cox and Cox, 2008) (Fig. 5). With the increase of ASL at a given *k*, the clusters become more separable (i.e., each row in Fig. 5 represents the three experiments with the same *k*).

**Figure 5.**
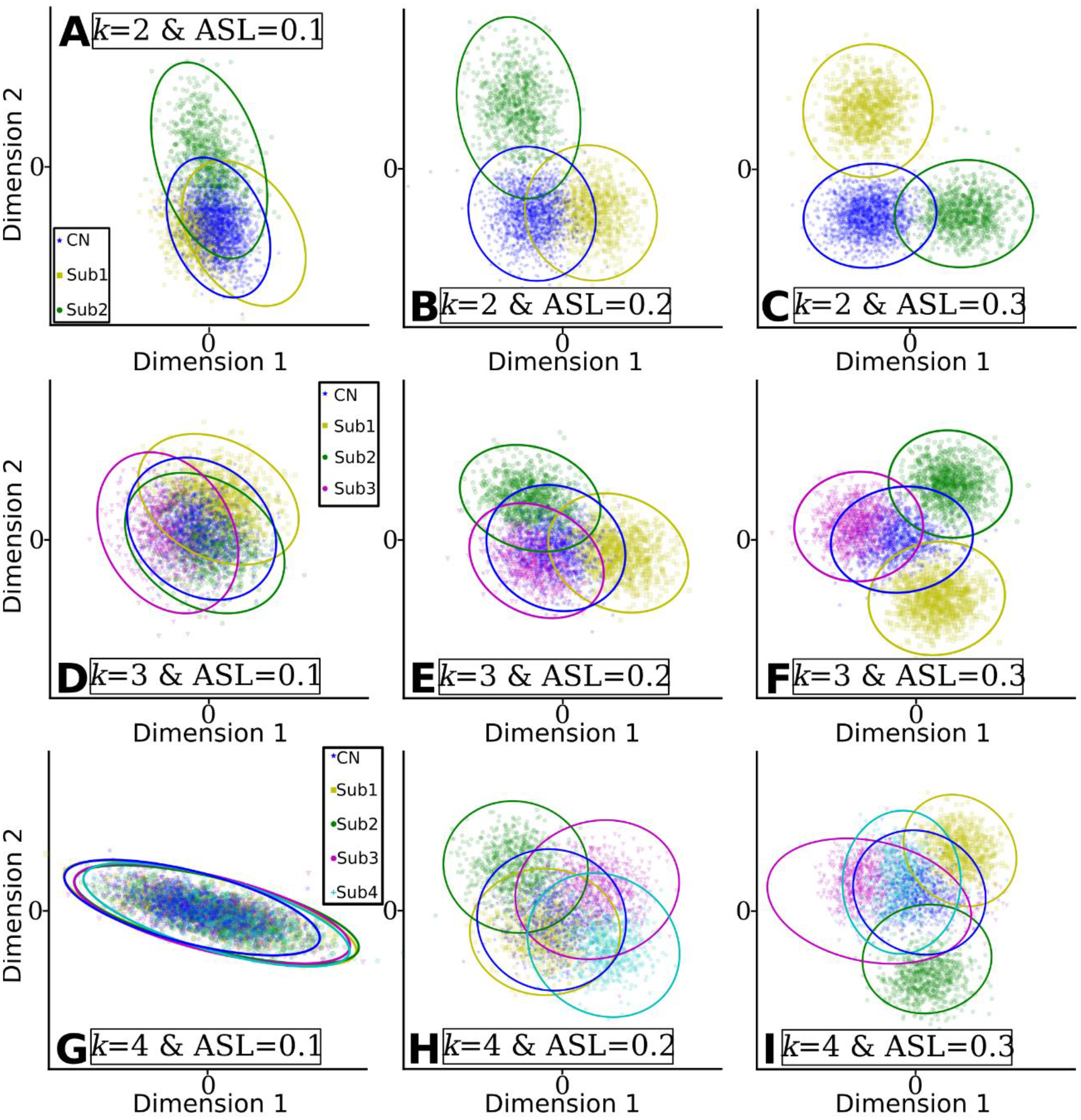
Clusters found by MAGIC become more distinguishable when the clustering conditions are favorable, i.e., higher ASL or lower *k*. The clusters were projected into 2D space for visualization. Dimension 1 and Dimension 2 represent the two components projected by multidimensional scaling methods. A) *k*=2 & ASL=0.1; B) *k*=2 & ASL=0.2; C) *k*=2 & ASL=0.3; D) *k*=3 & ASL=0.1; E) *k*=3 & ASL=0.2; F) *k*=3 & ASL=0.3; G) *k*=4 & ASL=0.1; H) *k*=4 & ASL=0.2; I) *k*=4 & ASL=0.3.

#### 4.2.5. Influence of overlapping atrophy patterns

We generated overlapping atrophy patterns based on the original experiments for each *k*. For *k*=2, Sub2 had subcortical atrophy in the initial experiments (Fig. 3), and we additionally simulated parietal atrophy. Similarly, for *k*=3 and 4, global cortical atrophy was imposed within Sub1 (frontal atrophy subtype in the original experiments) and Sub3 (temporal atrophy subtype in the initial experiments) members, respectively. The ground truth of overlapping neuroanatomical patterns is detailed in supplementary eFigure 2.

As shown in Table 3, MAGIC obtained inferior clustering performance compared to the original experiments for i) *k*=2 & ASL=0.1, ii) *k*=3 & ASL=0.1, iii) *k*=3 & ASL=0.2 and iv) *k*=4 & ASL=0.2, and comparable results for experiments with ASL=0.3. The results for the ARI_CV, voxel-wise effect size maps and the 2D visualization of subtypes are presented in supplementary eFigure 2, 3 and 4, respectively.

**Table 3.** Comparison of the original clustering performance (left column) to the influence of overlapping atrophy patterns (middle column) and the larger brain volume (right column). Compared to the original experiments, overlapping atrophy patterns result in lower clustering performance, while larger brain volume shows no extensive clustering performance effects.

| Experiment | Original experiments | Overlapping atrophy patterns | Larger brain volume |
| --- | --- | --- | --- |
| $k=2$ & ASL=0.1 | 0.610 | 0.501 | 0.562 |
| $k=2$ & ASL=0.2 | 0.960 | 0.946 | 0.947 |
| $k=2$ & ASL=0.3 | 0.962 | 0.962 | 0.962 |
| $k=3$ & ASL=0.1 | 0.368 | 0.281 | 0.393 |
| $k=3$ & ASL=0.2 | 0.934 | 0.879 | 0.926 |
| $k=3$ & ASL=0.3 | 0.995 | 0.977 | 0.976 |
| $k=4$ & ASL=0.1 | 0.091 | 0.111 | 0.210 |
| $k=4$ & ASL=0.2 | 0.713 | 0.628 | 0.731 |
| $k=4$ & ASL=0.3 | 0.966 | 0.967 | 0.965 |

#### 4.2.6. Influence of larger regional brain volumes

Instead of simulating brain atrophy as in the original experiments (Fig. 3), we introduced larger brain volumes by increasing the voxel’s intensity value inside the pre-defined patterns for Sub2 members for experiments *k*=2, Sub3 members for experiments *k*=3 and Sub4 members for experiments *k*=4. The simulated neuroanatomical patterns are detailed in supplementary eFigure 5.

As shown in Table 3, MAGIC obtained comparable clustering performance to all settings’ original experiments. The results for the ARI_CV, voxel-wise effect size maps, and the 2D visualization of subtypes are presented in supplementary eFigure 5, 6, and 7, respectively.

**Figure 6.**
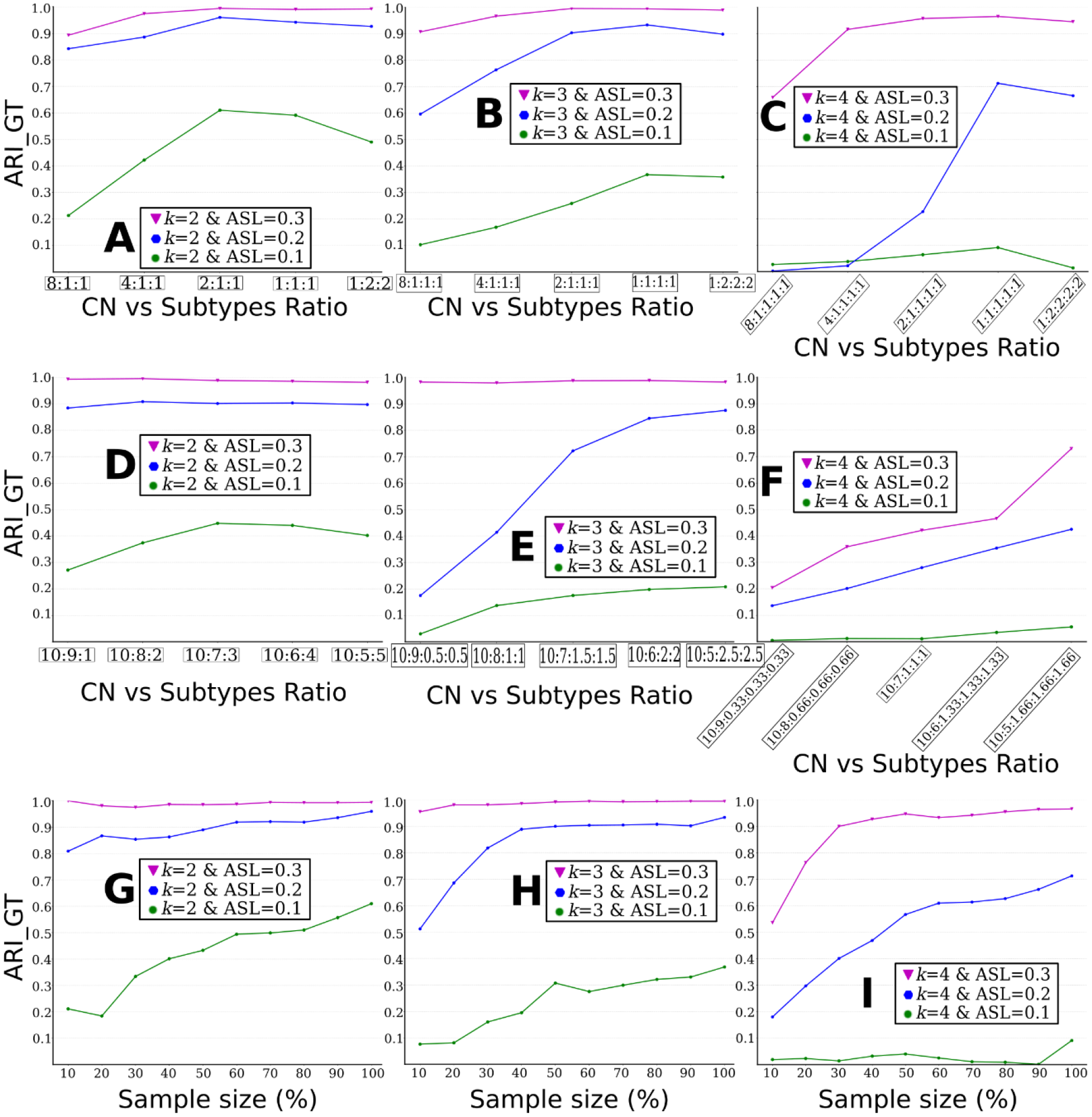
The influence of different ratios of imbalanced data between CN vs. subtypes is presented in Fig. A, B, and C, among subtypes in Fig. D, E and F. The influence of sample size is displayed in Fig. G, H, and I. A) influence of data imbalance between CN and subtypes for *k*=2; B) influence of data imbalance between CN and subtypes for *k*=3; C) influence of data imbalance between CN and subtypes for *k*=4; D) influence of data imbalance among subtypes for *k*=2. Clustering performance improves with the increase of the sample size. E) influence of data imbalance among subtypes for *k*=3; F) influence of data imbalance among subtypes for *k*=4; G) influence of sample sizes for *k*=2; H) influence of sample sizes for *k*=3; I) influence of sample sizes for *k*=4.

#### 4.2.7. Influence of data imbalance

We first evaluated the influence of data imbalance for CN vs. subtypes. The imbalance ratios were achieved by randomly subsampling from the groups of subtypes. As shown in Fig. 6 A, B, and C, clustering performance considerably increased when the groups became more balanced. With the highest imbalance ratio (8:1), all experiments obtained the lowest ARI_GTs. Generally, the ratios of 2:1 performed on par with the ratios of 1:1 and 1:2.

We then evaluated the influence of data imbalance among subtypes by assuming that CN and PT (sum of all subtypes) were balanced (Fig. 6D, E, and F). Similarly, clustering performance considerably increased with more balanced data. On the other hand, when ASL is large (i.e., 0.3), data imbalance showed a limited impact on clustering performance (e.g., Fig. 6D and E).

#### 4.2.8. Influence of sample size

The influence of the sample size on clustering performance was assessed (Fig. 6G, H and I). For each experiment, MAGIC was run with data ranging from 10% to 100% of the sample size by keeping the original group ratios unchanged (i.e., CN vs. Sub1 vs. Sub2 vs …).

Generally, clustering performance improved with the increasing sample size. For experiments *k*=2 & ASL=0.3 and *k*=3 & ASL=0.3, clustering performance was almost perfect at all different sample size choices. For experiment *k*=4 & ASL=0.1 (Fig. 6I), MAGIC obtained poor clustering performance.

### 4.3. Experiments using Alzheimer’s disease data

When applied to ADNI data, ARI_CV was the highest at *k*=2 (<0.5), compared to other values of *k* (Fig. 7A). For *k*=2, The effect size maps revealed two distinct neuroanatomical patterns: i) Sub1 (*N*=396) showed relatively normal brain anatomy, except for focalized brain atrophy in subcortical regions. In contrast, Sub2 (*N*=791) had diffuse atrophy with the largest effect size (Cohen’s *f*2 = 0.45) in the hippocampus, amygdala, and temporal regions (Fig. 7B). For *k*=3, the three subtypes all presented diffuse brain atrophy (Fig. 7C). For *k*=4, Sub1 (*N*=363) showed only focal atrophy in temporal regions. Sub2 (*N*=416) is the typical AD pattern showing whole-brain atrophy and most severe atrophy in temporal and hippocampus regions. Sub3 (*N*=210) showed atypical AD patterns without affecting the hippocampus and temporal lobes (Fig. 7D). Sub4 (*N*=198) preserved relatively normal brain anatomy. Large effect sizes were detected in all subtypes but the neuroanatomical patterns overlapped and were focalized.

**Figure 7.**
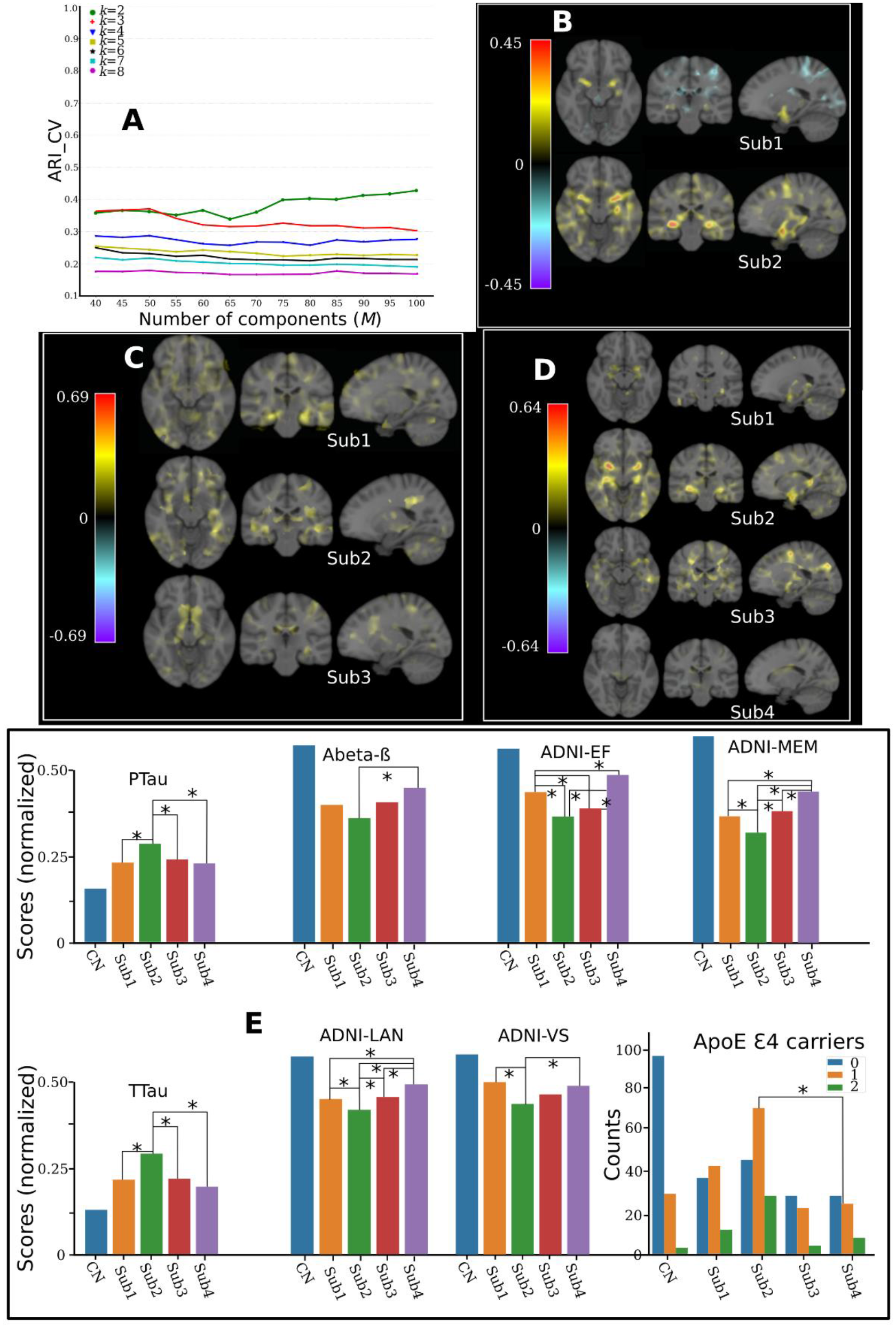
Applying MAGIC to AD and MCI patients of ADNI. A) Cross-validation for choosing the “optimal” *k*. In general, the ARI_CV is low for all different choices of *k* and with the highest for *k*=2. B) Voxel-wise effect size (Cohen’s *f*2) maps for the neuroanatomical patterns between the two subtypes and CN. C) Effect size maps for the three subtypes and CN neuroanatomical patterns. D) Effect size maps for the neuroanatomical patterns between the four subtypes and CN. E) Clinical characteristics for the four subtypes and CN. Mann–Whitney– Wilcoxon test was used for continuous variables and the Chi-square test of independence for categorical variables. The significance threshold is 0.05 and * denotes statistical significance. All continuous variables were normalized for visualization purposes. We harmonized the low-dimensional components using the Combat-GAM model (Pomponio et al., 2019) to mitigate the site effect.

The CV procedure obtained consistently higher ARI_CV for *k*=2. It generally divides the patients into mild and severe atrophied groups, which might not be clinically interesting. Using different semi-supervised clustering techniques but similar populations (AD and MCI from ADNI), we previously found four distinct subtypes (Dong et al., 2016b; Yang et al., 2021). Moreover, these subtypes have been previously reported in the works from other groups (see the Discussion section). To sum up, the results of ARI_CV, together with our semi-simulated experiments (Fig. 2D), might indicate that the CV procedure does not detect the true *k* due to unfavorable clustering conditions (e.g., the focalized effects or small sample size). Taken all together, we focused on *k*=4 for subsequent ADNI analyses.

To support our claims, we compared the clinical characteristics of the four subtypes (Fig. 7E). Details are presented in supplementary eTable 1 for statistics and data availability. In general, Sub2 showed the highest TTau (127.62) and PTau (45.16), the highest ApoE Ɛ4 carrier rate (68%), and the most deficient cognitive performance across the four domains, whereas Sub4 showed the opposite trend and represented a more normal-like clinical and neuroimaging profile.

We demonstrated the utility of MAGIC for a binary classification task (541 CN vs 339 AD) using ADNI data and compared it to HYDRA and a linear SVM. We adopted the same cross-validation (CV) procedure as MAGIC for all models for a fair comparison. The constructed polytope was used for classification with MAGIC and HYDRA. MAGIC (0.85±0.03) obtained slightly better performance than HYDRA (0.84±0.04) and a linear SVM (0.82±0.04) (supplementary eFigure 8).

**Figure 8.**
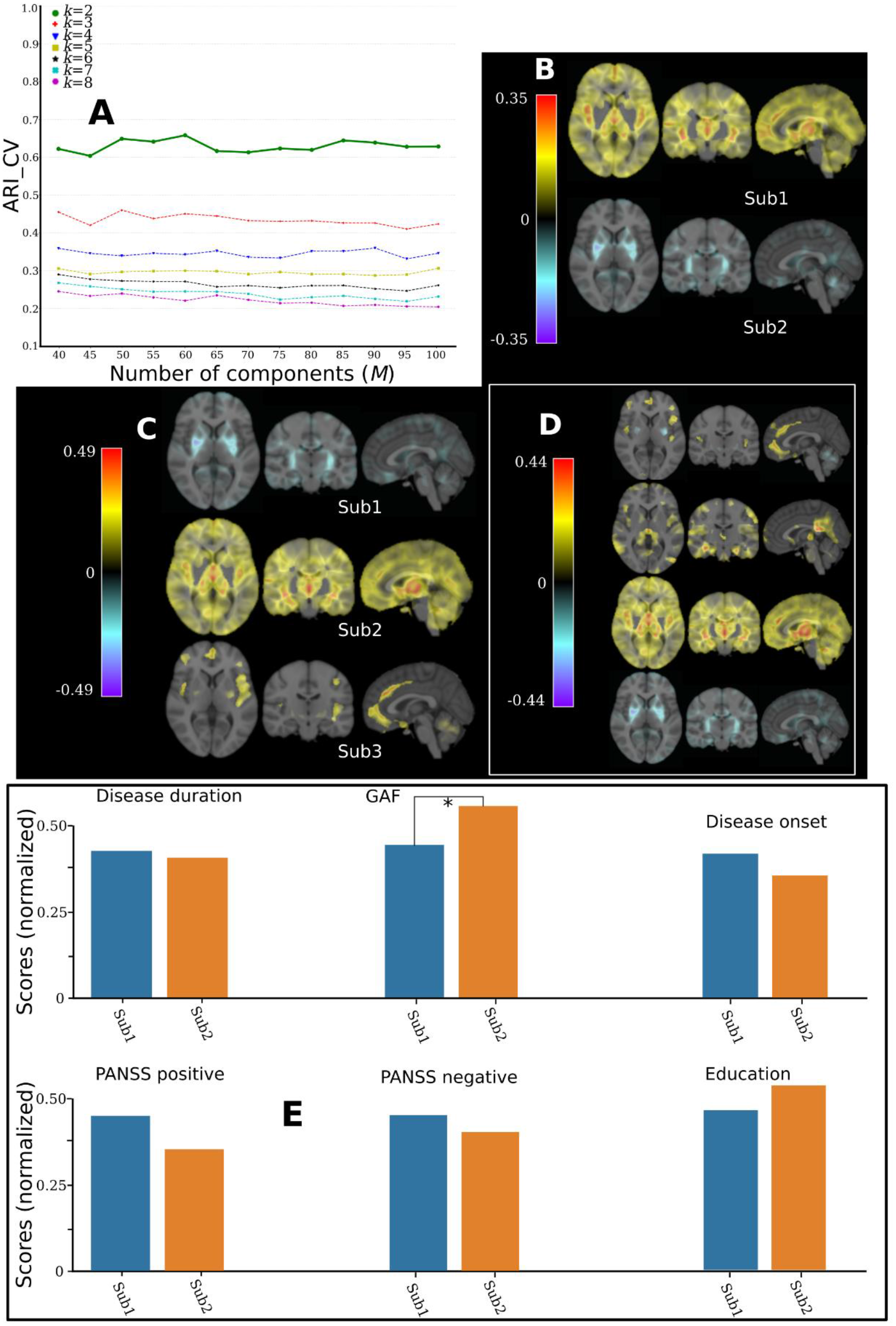
Applying MAGIC to schizophrenia patients from PHENOM. A) Cross-validation for choosing the “optimal” *k*. B) Voxel-wise effect size maps for the neuroanatomical patterns between the two subtypes and CN. C) Effect size maps for the neuroanatomical patterns between the three subtypes and CN. D) Effect size maps for the neuroanatomical patterns between the four subtypes and CN. E) Clinical characteristics for the two subtypes. Mann–Whitney–Wilcoxon test was used for continuous. The significance threshold is 0.05 and * denotes statistical significance. All continuous variables were normalized for visualization purposes. GAF: global assessment of functioning. We harmonized the low-dimensional components using the Combat-GAM model (Pomponio et al., 2019) to mitigate the site effect. Disease onset: age at onset.

### 4.4. Experiments using schizophrenia data

We then applied MAGIC to PHENOM data. For model selection, the CV procedure showed consistently higher ARI_CV values for *k*=2 (ARI_CV>0.6) compared to other resolutions (Fig. 8A). For *k*=2, the effect size maps revealed two distinct neuroanatomical patterns: Sub1 (*N*=383) showed widespread brain atrophy compared to CN, with the highest effect size (Cohen’s *f*2=0.35) in the thalamus and insula. In contrast, Sub2 (*N*=200) exhibited no brain atrophy but larger brain volumes than CN in the pallidum and putamen (Fig. 8B). For *k*=3, the first two subtypes [Sub1 (*N*=302) and Sub2 (*N*=185)] persisted, and Sub3 (*N*=96) showed atrophy in frontal lobe and insula regions (Fig. 8C). For *k*=4, Sub4 (*N*=109) showed diffuse atrophy patterns with a small effect size (Cohen’s *f*2<0.07) (Fig. 8D).

For *k*=2, MAGIC validated the two subtypes revealed in our previous work, in which HYDRA and a smaller sample size (364 CN and 307 SCZ) were used to derive the two subtypes (Chand et al., 2020). Subsequently, we compared the clinical characteristics of the two subtypes (Fig. 8E). Details are presented in supplementary eTable 2. Sub2 showed better performance in the global assessment of functioning (GAF) scale.

## 5. Discussion

### Synopsis

This paper presents MAGIC, a novel multi-scale semi-supervised clustering method for dissecting disease heterogeneity. The proposed method seamlessly integrates multi-scale representation learning and semi-supervised clustering in a coherent framework via a double-cyclic optimization procedure to yield scale-agnostic delineation of heterogeneous disease patterns. In contrast to existing unsupervised approaches presented in (Ezzati et al., 2020, 2020; Jeon et al., 2019, 2019; Jung et al., 2016; Lubeiro et al., 2016; Nettiksimmons et al., 2014; Ota et al., 2016; Pan et al., 2020; Park et al., 2017; Planchuelo-Gómez et al., 2020; Poulakis et al., 2020, 2020, 2018; Sugihara et al., 2016; Ten Kate et al., 2018), MAGIC is a semi-supervised approach, leveraging the patient-control dichotomy to drive subtypes that reflect distinct pathological processes. In contrast to the existing state-of-the-art semi-supervised clustering method (i.e., HYDRA), MAGIC can accurately delineate effect patterns that are both global and focal, thanks to its multi-scale optimization routine. The validity of MAGIC is demonstrated in semi-simulated experiments. We show MAGIC’s ability to discern disease subtypes and their neuroanatomical patterns under various simulated scenarios constructed by varying the ASL, sample size, and sample imbalance, respectively. More importantly, we showcased that the results of the analyses on ADNI and PHENOM datasets and the semi-simulated experiments exquisitely echo each other. Two clinically distinct subtypes of SCZ patients are established with high confidence, corresponding to the simulation where the clustering conditions are favorable. In contrast, the four subtypes in AD and MCI are less convincing, reflecting the simulation where the conditions are hard to disentangle.

### MAGIC outperforms comparable heterogeneity analysis methods

Concerning clustering performance, MAGIC outperformed competing methods. On the one hand, compared to HYDRA, the minor gain in clustering accuracy in MAGIC is likely driven by multi-scale features that can better explain the variance due to heterogeneity. We hypothesize that the opNMF multi-scale components accurately reflect multi-scale brain organization that has previously been demonstrated in network analysis (Betzel and Bassett, 2017), brain modeling (Schirner et al., 2018), and signal processing (Starck et al., 1998) in the literature. Furthermore, multi-scale learning has shown great potential in medical imaging for different tasks, such as segmentation (Doshi et al., 2016; Kamnitsas et al., 2017) or classification (Cui et al., 2016; Hu et al., 2016). On the other hand, MAGIC substantially outperforms unsupervised clustering methods. Since unsupervised clustering methods directly partition patient samples into clusters based on similarity/dissimilarity or distance (Altman and Krzywinski, 2017), they may be more likely driven by confounding factors such as brain size, age, and sex instead of pathology-related variations, which is partially addressed by MAGIC. Namely, MAGIC can derive pathology-driven subtypes in a multi-scale manner by leveraging the reference label (i.e., CN) and the fuzzy patient labels (i.e., PT).

### Under what conditions does MAGIC succeed or fail?

The critical yet challenging choice to be made in all algorithms related to clustering is to choose the appropriate number of clusters (Climescu-Haulica, 2007; Fu and Perry, 2020; Mirkin, 2011), since all clustering methods find patterns in data - whether they are real or not (Altman and Krzywinski, 2017). In addition to providing a new clustering method, we provided guidelines to these heterogeneity analysis algorithms’ practitioners. Specifically, our experiments shed light on selecting the number of clusters and provide criteria when the clustering analysis is reliable and when it needs to be approached with caution. In general, we suggest performing model selection using a cross-validation strategy based on clustering stability. In our experiments, ARI reliably recovered the ground truth number of clusters when the ASL and sample size were large and data was reasonably balanced. However, one should note that a lower number of clusters intrinsically gives more stable results (i.e., higher ARI_CV). In such cases, in actual clinical applications, other insights may be required to support further the subtypes found, such as the effect size map or prior clinical knowledge.

Different choices of the key components (e.g., sample size or data imbalance) have detrimental or positive influences on clustering. With the increase of the complexity of clustering (e.g., increasing the number of clusters or decreasing ASL), MAGIC’s clustering performance degrades gradually. This is to be expected as the boundaries between clusters become increasingly blurred and indivisible. Moreover, imbalanced data have adverse effects on clustering results. This is in line with previous findings (Dubey et al., 2014; Samper-González et al., 2018). The authors found that balanced data obtained better classification results than the imbalanced using T1w MRI from ADNI. Note that MAGIC essentially performs clustering and supervised classification simultaneously. Unsurprisingly, increased sample size leads to better clustering performance, consistent with the findings from previous studies (Abdulkadir et al., 2011; Chu et al., 2011; Franke et al., 2010; Samper-González et al., 2018; Schulz et al., 2020b). The rule of thumb in semi-supervised clustering is to collect moderately balanced data and large samples in practice. Finally, instead of larger brain patterns, adding overlapping patterns made it more difficult to disentangle these effects. The former situation is more commonly present in practice. In these cases, larger sample cohorts may be needed to unravel the overlapping heterogeneity.

Our semi-simulated experiments are of great value for assessing clustering results. They enable us to understand potential false-positive results. Namely, suppose the sample size, ARI_CV, and/or the effect sizes are low. In that case, none of the existing methods may uncover the true heterogeneity or may reveal a lower number of subtypes than what is present.

### Subtypes in AD and MCI

Applying the proposed method to structural imaging data from ADNI indicates that the current sample size or technique cannot detect the “true” number of clusters. This result is consistent with the semi-simulated experiment, in which the resolution of *k*=2 intrinsically tends to be higher when the clustering conditions are less favorable (Fig. 2D). In such cases, clinical priors and external validation of the claimed subtypes are highly demanded. Furthermore, neurodegenerative diseases are highly heterogeneous, and the neuroanatomical patterns might be highly overlapping. For instance, previous autopsy studies (DeTure and Dickson, 2019; Perl, 2010) have shown that pure AD cases are relatively infrequent, as AD and other comorbid conditions (e.g., vascular disease or Lewy body disease) often co-exist (Rabinovici et al., 2016).

Among the four subtypes, Sub4 displays normal brain anatomy (Fig. 7D). This was supported by the distribution of AD/MCI. Sub4 (164 MCI & 34 AD) has the highest proportion of MCI. The normal anatomy subtype has been confirmed in previous works both from semi-supervised and unsupervised methods (Dong et al., 2016b; Ezzati et al., 2020; Jung et al., 2016; Nettiksimmons et al., 2014; Ota et al., 2016; Poulakis et al., 2020, 2018; Ten Kate et al., 2018; Yang et al., 2020). Sub2 showed typical AD-like neuroanatomical patterns with diffuse atrophy over the whole brain, with the largest effect size in the hippocampus and medial temporal lobe. Those affected regions have been widely reported as hallmarks of AD in case-control studies (Hanyu et al., 1998; Müller et al., 2005; Varghese et al., 2013) and have been confirmed in previous clustering literature (Dong et al., 2016b; Nettiksimmons et al., 2014; Noh et al., 2014; Poulakis et al., 2018; Ten Kate et al., 2018; Varol et al., 2017; Yang et al., 2020; Young et al., 2018). Conversely, Sub3 showed an atypical widespread atrophy pattern that did not include the hippocampus and the temporal lobe (Dong et al., 2016b; Poulakis et al., 2018; Yang et al., 2020).

The clinical characteristics of the four subtypes are in line with their neuroanatomical patterns (Fig. 7E). The normal-like subtype had the lowest level with respect to CSF amyloid-b 1-42, CSF-tau levels, most minor cognitive impairment. Moreover, despite the methodological differences across studies, the resulting subtypes’ agreement emphasizes that AD should be considered a neuroanatomically heterogeneous disease. These distinct imaging signatures or dimensions may elucidate different brain mechanisms and pathways leading to AD and eventually contribute to the refinement of the “N” dimension in the “A/T/N” system (Jack et al., 2016).

### Subtypes in schizophrenia

MAGIC discovered two highly reproducible and neuroanatomically distinct subtypes in schizophrenia patients. MAGIC obtained consistently higher clustering stability for *k*=2 (ARI_CV>0.6) and large effect size for subtypes’ neuroanatomical patterns. Furthermore, these two subtypes were retained for a higher resolution of the number of subtypes (*k*>2). Sub1’s neuroanatomical patterns are in line with previous case-control literature, demonstrating widespread GM atrophy (Okada, N et al., 2016; Rozycki et al., 2018; van Erp et al., 2016). In contrast, Sub2 showing larger brain volume in basal ganglia corresponds to previous works reporting subcortical GM increases (Brugger and Howes, 2017; Okada, N et al., 2016, 2016; W. Zhang et al., 2015). Moreover, Sub2 showed less functional disability (higher GAF). The two subtypes found by MAGIC reinforce the need for the refinement of the neuroanatomical dimensions in schizophrenia.

### Potential and challenges

The application of clustering methods to neuroimaging data has recently drawn significant attention and has led to several key publications in recent years. Herein we demonstrated MAGIC’s potential for dissecting the neuroanatomical heterogeneity of brain diseases, indicating that the current “all-in-one-bucket” diagnostic criteria may not be appropriate for certain neurodegenerative and neuropsychiatric disorders. On the other hand, clustering methods always end up with clusters, even if there are no natural clusters in the data (Altman and Krzywinski, 2017). If they indeed exist, the disease subtypes often present neuroanatomically overlapping patterns, unlike the semi-simulated conditions with purely defined orthogonal patterns. None of the heterogeneity analysis tools were sufficiently powered to accurately disentangle heterogeneity in small sample cohorts or with weak discriminative power of pattern identifiability in our experiments. In this case, care must be taken to provide additional information, such as external validation of subtypes to clinical profiles, to substantiate any clinical interpretation of the identified subtypes.

To sum up, our semi-simulated results and the application to ADNI and PHENOM datasets emphasize the value of the current work. We provide the semi-simulated experiments and the proposed model to the community and offer new vistas for future research in refining subtypes in brain diseases. The reproducibility of clustering, effect size maps of subtypes, and sample imbalance should be carefully examined. Ultimately, good practices, such as extensive reproducibility analyses, including permutation tests (Chand et al., 2020), should be performed to support the subtypes’ stability and robustness. However, we observed a steady improvement of clustering performance with increased sample sizes even with overlapping anatomical patterns. This is a promising sign for the utility of these machine learning-based clustering tools with the increasing demand for large neuroimaging consortia.

Our model nevertheless has the following limitations. First, MAGIC is designed for “pure” clustering tasks that seek the disease’s subtypes without considering the disease progression factors or stages (Young et al., 2018). A future direction is extending MAGIC to assign subtypes to longitudinal scans and study disease progression. Moreover, the sample size of AD necessary to draw a solid conclusion for those subtypes may be larger than analyzed. Clustering performance was positively associated with sample size in our simulation. Moreover, a possible extension of the proposed method is integrating clinical or genetic data to derive subtypes that show consistency across different modalities. Lastly, external validation of the claimed AD subtypes to an independent dataset, such as the Dementias Platform UK (DPUK) (Bauermeister et al., 2020), is another future direction.

## Table of Abbreviations

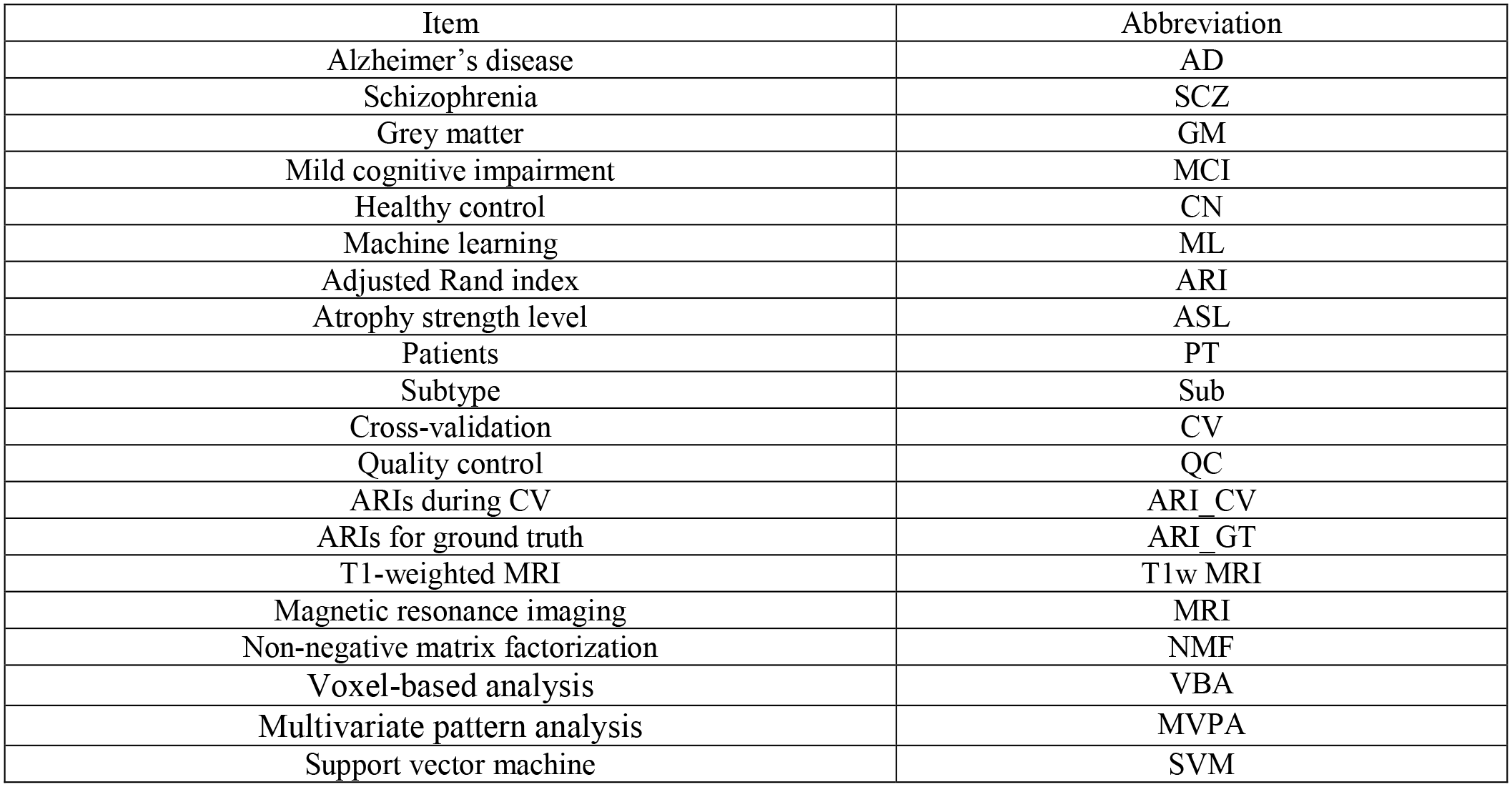

## Table of variables

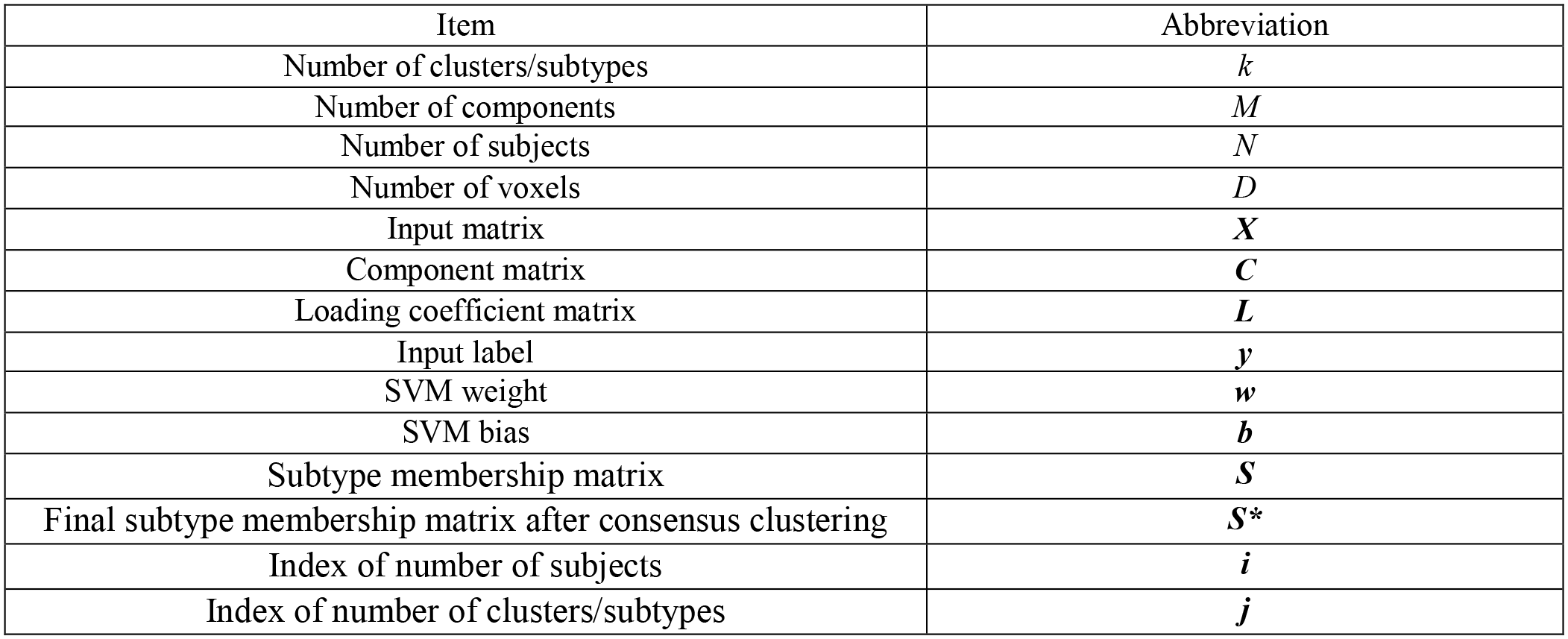

## Acknowledgements

This work was granted access to be supported in part by NIH grants R01MH112070, 1U01AG068057, 1RF1AG054409, and R01AG067103. Additional support was provided by S10OD023495, R01MH101111, R01MH113565, R01MH113550, R01MH112847, and R01EB022573. The work was also supported by the PRONIA project as funded by the European Union 7th Framework Program grant 602152. Data collection and sharing for this project were funded by the Alzheimer’s Disease Neuroimaging Initiative (ADNI) (National Institutes of Health Grant U01 AG024904) and DOD ADNI (Department of Defense award number W81XWH-12-2-0012). ADNI is funded by the National Institute on Aging, the National Institute of Biomedical Imaging and Bioengineering, and through generous contributions from the following: AbbVie, Alzheimer’s Association; Alzheimer’s Drug Discovery Foundation; Araclon Biotech; BioClinica, Inc.; Biogen; Bristol-Myers Squibb Company; CereSpir, Inc.; Cogstate; Eisai Inc.; Elan Pharmaceuticals, Inc.; Eli Lilly and Company; EuroImmun; F. Hoffmann-La Roche Ltd and its affiliated company Genentech, Inc.; Fujirebio; GE Healthcare; IXICO Ltd.; Janssen Alzheimer Immunotherapy Research & Development, LLC.; Johnson & Johnson Pharmaceutical Research & Development LLC.; Lumosity; Lundbeck; Merck & Co., Inc.; Meso Scale Diagnostics, LLC.; NeuroRx Research; Neurotrack Technologies; Novartis Pharmaceuticals Corporation; Pfizer Inc.; Piramal Imaging; Servier; Takeda Pharmaceutical Company; and Transition Therapeutics. The Canadian Institutes of Health Research is providing funds to support ADNI clinical sites in Canada. Private sector contributions are facilitated by the Foundation for the National Institutes of Health (www.fnih.org). The grantee organization is the Northern California Institute for Research and Education, and the study is coordinated by the Alzheimer’s Therapeutic Research Institute at the University of Southern California. ADNI data are disseminated by the Laboratory for Neuro Imaging at the University of Southern California. This research has also been conducted using the UK Biobank Resource (UKBB Application Number: 35148): https://www.ukbiobank.ac.uk/.

## eMethod 1. HYDRA

The core motor of the HYDRA algorithm is the non-linear polytope that is constructed by multiple (number of clusters) linear hyperplanes from linear SVMs. Herein, we briefly introduce the mathematical fundamentals of HYDRA. For more details, we encourage the readers to refer to (Varol et al., 2017).

### Support vector machine

SVM aims to estimate a hyperplane that separates the two groups by a half-space while ensuring that the distance/margin from the hyperplane is maximized for each data point. Let us denote ***w*** as all possible linear classifiers in the set *Ƒ* for a given dataset *Ɗ*. The goal of an SVM is to find the classifier belonging to the set *Ƒ* that maximizes the margin:

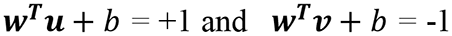

where the set of points from both groups ***u*** and ***v*** satisfy the equation. The margin can be derived as 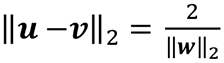, since ***u*** and ***v*** are parallel with each other.

This leads to the well-known SVM objective:

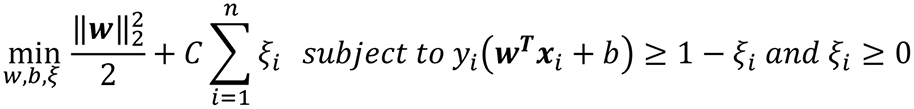

where *ξ*_*i*_ represents the slack term when the two groups are non-separable.

### Convex polytope construction via linear SVMs

HYDRA exquisitely constructs a “polytope” that can separate each cluster/subtype with a face of the polytope – that is, derived from a linear SVM. Let’s confine the control group to its interior region of the polytope and the patient group to its exterior. Thus, the search space Ƒ_*k*_ can be defined as:

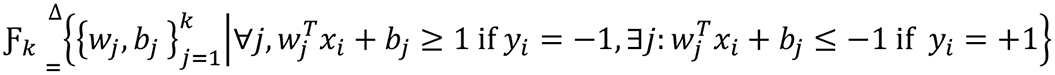

In short, the set Ƒ_*k*_ comprises all sets of *k* linear SVMs such that all CN subjects were correctly classified, while every patient was at least precisely classified by one hyperplane. Subsequently, deriving the clustering membership is straightforward. The search space can be rewritten as:

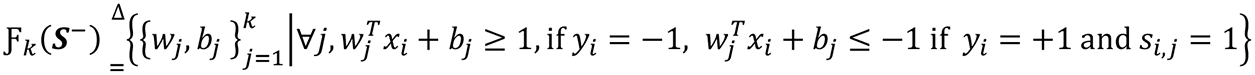

where ***S***^−^ ∈ {0, 1}^*n*−x*k*^ denotes the binary matrix that describes the assignment of the *i*-th patient sample to the *j*-th face of the polytope. HYDRA maximizes the average margin across all faces of the polytope. The objective of HYDRA is separable into *k*-independent subproblems; each subproblem is analogous to the SVM formulation:

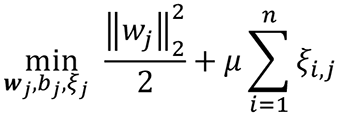

Where *μ* is the penalty parameter on the training error. If we consider all *k*-hyperplanes together, this leads to the Equation. 3 in the main manuscript.

## eMethod 2. Main datasets

Three datasets have been mainly used for the current study: the UK Biobank (UKBB) studies, the Alzheimer’s Disease Neuroimaging Initiative (ADNI), and the Psychosis Heterogeneity Evaluated via Dimensional Neuroimaging (PHENOM) study. In the following, we briefly describe these datasets.

### ● UK Biobank (UKBB)

For our study sample, we used the UK Biobank, a dataset of approximately 500,000 UK adults sampled via population-based registries (http://www.ukbiobank.ac.uk). The UK Biobank received ethical approval from the National Research Ethics Service Committee North West–Haydock (reference 11/NW/0382). All participants provided informed consent and were aged from approximately 40 to 69 years of age at the time of enrollment (http://biobank.ctsu.ox.ac.uk/crystal/field.cgi?id=200). Participants were recruited from across the United Kingdom, and initial enrollment was carried out from 2006 to 2010. Participants provided socio-demographic, cognitive, and medical data via questionnaires and physical assessments. Starting in 2014, a subset of the original sample later underwent magnetic-resonance brain imaging (MRI) (UK Biobank Brain Imaging Documentation, http://www.ukbiobank.ac.uk). The MRI data used in the current study were acquired between 2014 and 2019. Specifically, in our semi-simulated experiments, we finally included 4403 subjects whose T1-weighted (T1w) MRI was available (UKBB Application Number 35148).

### ● Alzheimer’s Disease Neuroimaging Initiative (ADNI)

Part of the data used in the preparation of this article was obtained from the Alzheimer’s Disease Neuroimaging Initiative (ADNI) database (adni.loni.usc.edu). The ADNI was launched in 2003 as a public-private partnership, led by Principal Investigator Michael W. Weiner, MD. The primary goal of ADNI has been to test whether serial magnetic resonance imaging (MRI), positron emission tomography (PET), other biological markers, and clinical and neuropsychological assessment can be combined to measure the progression of mild cognitive impairment (MCI) and early Alzheimer’s disease (AD). For up-to-date information, see www.adni-info.org.1 The ADNI study is composed of 4 cohorts: ADNI-1, ADNI-GO, ADNI-2, and ADNI-3. These cohorts are dependent and longitudinal, meaning that one cohort may include the same patient more than once and that different cohorts may include the same patients. Diagnosis labels are given by a physician after a series of tests (Petersen et al., 2010). The existing labels are:

- AD (Alzheimer’s disease): mildly demented patients,
- MCI (mild cognitive impairment): patients in the prodromal phase of AD,
- NC (normal controls): elderly control participants,
- SMC (significant memory concern): participants with cognitive complaints and no pathological neuropsychological findings. The designations SMC and subjective cognitive decline (SCD) are equivalently found in the literature.

Since the ADNI-GO and ADNI-2 cohorts, new patients at the very beginning of the prodromal stage have been recruited (Aisen et al., 2010), hence the MCI label has been split into two labels:

- EMCI (early MCI): patients at the beginning of the prodromal phase,
- LMCI (late MCI): patients at the end of the prodromal phase (similar to the previous label MCI of ADNI-1).

This division is made on the basis of the score obtained on memory tasks corrected by the education level. However, both classes remain very similar, and they are fused in many studies under the MCI label. We downloaded ADNI 1 and 2 data in December 2017, then added a small set of ADNI 3 in June 2020 for this work.

### ● Psychosis Heterogeneity Evaluated via Dimensional Neuroimaging (PHENOM)

Progress in delineating schizophrenia brain subtypes requires increased sample sizes, increased sample heterogeneity, and methodological advances that generalize across disparate sites and ethnicities. To respond to this challenge, we established a consortium spanning three continents called PHENOM (’Psychosis Heterogeneity Evaluated via Dimensional Neuroimaging’) (Chand et al., 2020; Satterthwaite et al., 2010; Wolf et al., 2014; T. Zhang et al., 2015; Zhu et al., 2016; Zhuo et al., 2016). We included chronic schizophrenia patients from eight international cohorts in the current study. The PHENOM sample comprises individuals with established schizophrenia (N=583) and healthy controls (N=583), including data from the USA, Germany, China, Australia, and Netherland. In the USA, subjects were recruited at the University of Pennsylvania and provided written informed consent under a protocol approved by the Institutional Review Board. Expert clinicians conducted the subject assessment. Diagnostic assessment employed the Structured Clinical Interview for DSM-IV (SCID). Subject exclusion criteria were a history of substance abuse in the past six months or a positive urine drug screen on the day of the study. Healthy control subjects were excluded if they met any DSM-IV psychiatric disorder criteria. Scale for the Assessment of Positive Symptoms (SAPS) and the Scale for the Assessment of Negative Symptoms (SANS) was assessed for patients. In Germany, subjects were recruited at Ludwig-Maximilians University following the ethics committee’s study approval. Subjects provided their written informed consent. Patient assessments were carried out by expert clinicians. The assessment included the SCID for Axis I & II disorders (SCID-I/-II), a clinical semi-standardized evaluation of medical and psychiatric history, review of medical records and psychotropic medications, and the evaluation of the Positive and Negative Syndrome Scale (PANSS) for disease severity and psychopathology. Individuals were excluded if they had other psychiatric and/or neurological diseases, past or present regular alcohol abuse, consumption of illicit drugs, past head trauma with loss of consciousness or electroconvulsive treatment, insufficient knowledge of German, IQ < 70, and age < 18 or > 65 years. Healthy controls with a positive familial history for mental illnesses (first-degree relatives) were also excluded. In China, subjects were recruited at Tianjin Medical University General Hospital following the Ethics Committee’s study approval. Each subject provided written informed consent. Diagnosis of patients was assessed following two clinical psychiatrists’ consensus using DSM-IV/SCID. Inclusion criteria were 16–60 years of age and right-handedness. Exclusion criteria were MRI contraindications, pregnancy, histories of systemic medical illness, central nervous system disorder and head trauma, and substance abuse within the last three months or lifetime history of substance abuse or dependence. The exclusion criteria were a history of psychiatric disease and first-degree relatives with a psychotic disorder for healthy control subjects. PANSS scores were assessed for patients for disease severity and psychopathology.

---

**eFigure 1.**
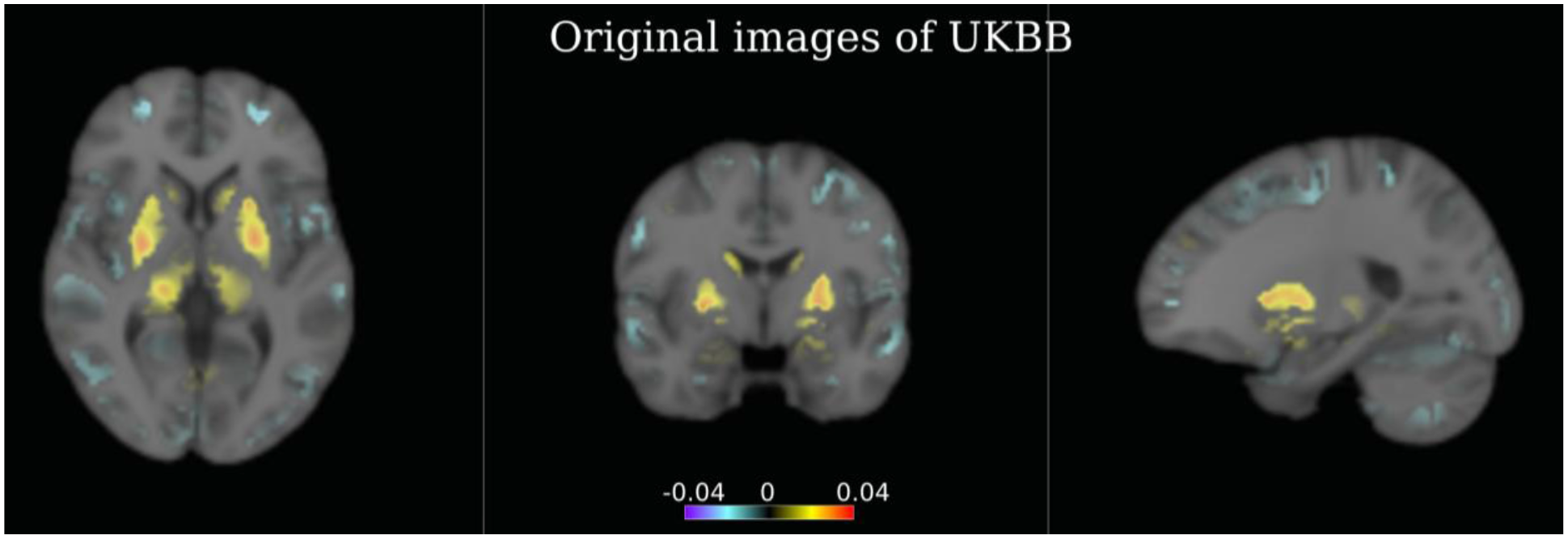
Voxel-wise effect size maps for original UKBB subjects from Sub1 and healthy control group before any simulation. Abnormality patterns exist in healthy control populations caused by typical brain aging. Therefore, the heterogeneity the proposed method recovered includes two sources: i) the simulated effect to mimic the disease effects and ii) the heterogeneity caused by typical brain aging in the original voxels.

**eFigure 2.**
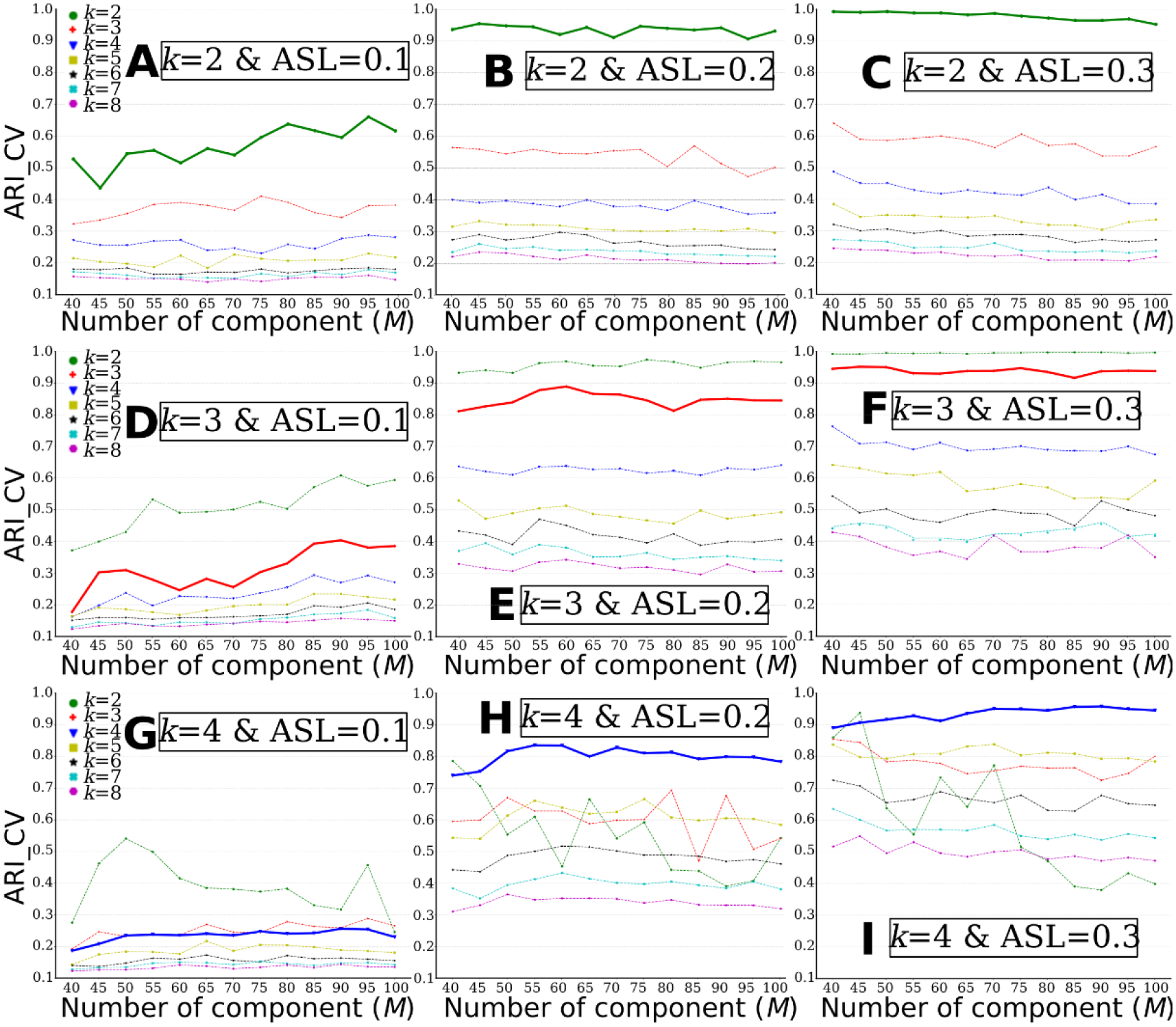
MAGIC finds the ground truth of the number of clusters (*k*) when the clustering conditions are favorable, i.e., higher ASL or lower *k*. The “optimal” k was determined based on the ARI (ARI_CV). A) k=2 & ASL=0.1; B) k=2 & ASL=0.2; C) k=2 & ASL=0.3; D) k=3 & ASL=0.1; E) k=3 & ASL=0.2; F) k=3 & ASL=0.3; G) *k*=4 & ASL=0.1; H) *k*=4 & ASL=0.2; I) *k*=4 & ASL=0.3. The bold lines represent the ground truth of *k* for each experiment.

**eFigure 3.**
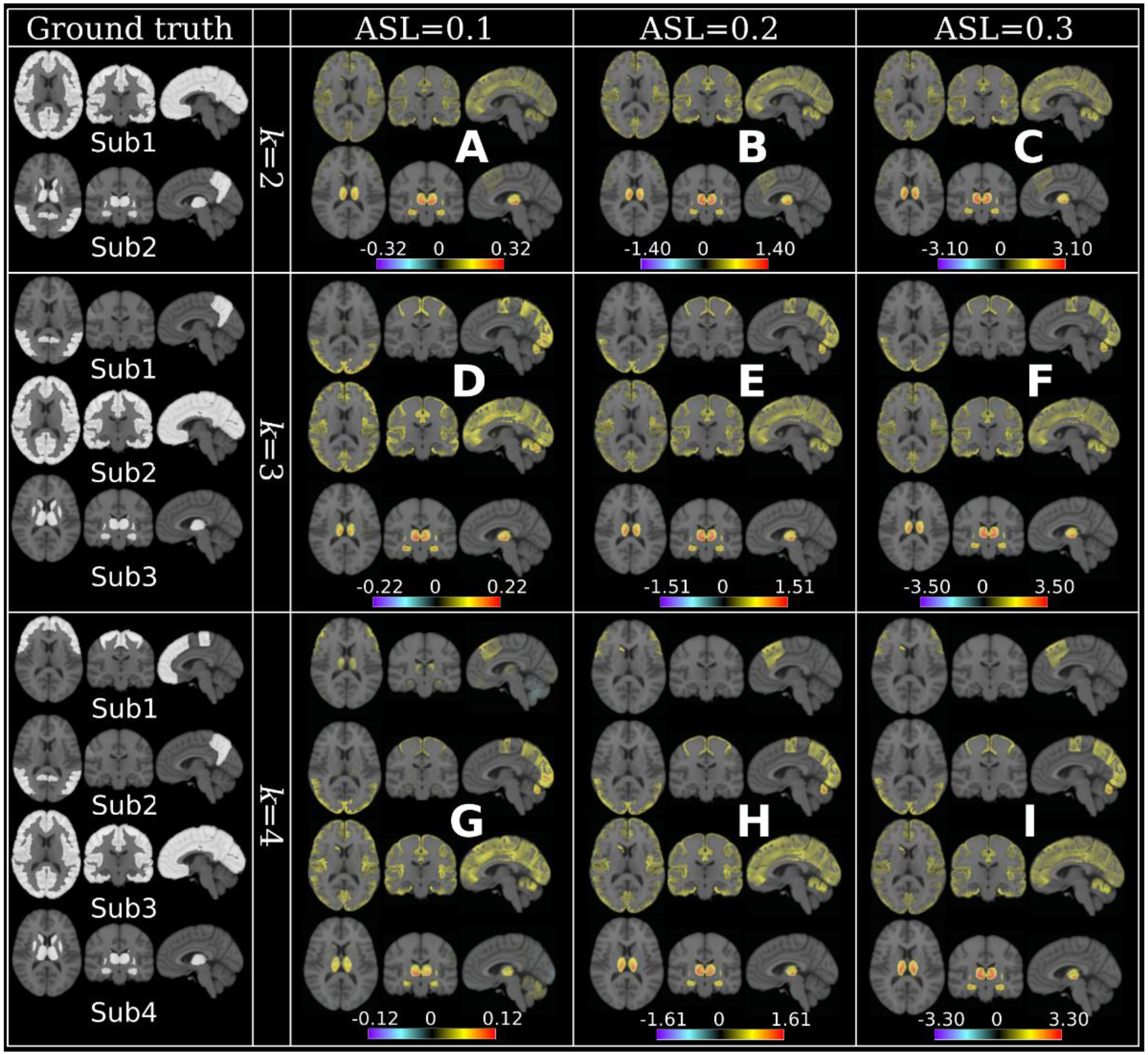
MAGIC finds the ground truth of subtype’s neuroanatomical patterns when the clustering conditions are favorable, i.e., higher ASL or lower k. Neuroanatomical patterns are displayed using effect size maps based on voxel-wise group comparisons between CN and subtypes. Positive values denote brain atrophy (CN > Sub), while negative values correspond to larger brain volume in subtypes (CN < Sub). The ground truth of the subtypes pattern is presented with a binary mask (white) for each k in the first column. A) k=2 & ASL=0.1; B) k=2 & ASL=0.2; C) k=2 & ASL=0.3; D) k=3 & ASL=0.1; E) k=3 & ASL=0.2; F) k=3 & ASL=0.3; G) k=4 & ASL=0.1; H) k=4 & ASL=0.2; I) k=4 & ASL=0.3. For reference, Cohen’s f2 ≥ 0.02, ≥ 0.15, and ≥ 0.35 represent small, medium, and large effect sizes, respectively.

**eFigure 4.**
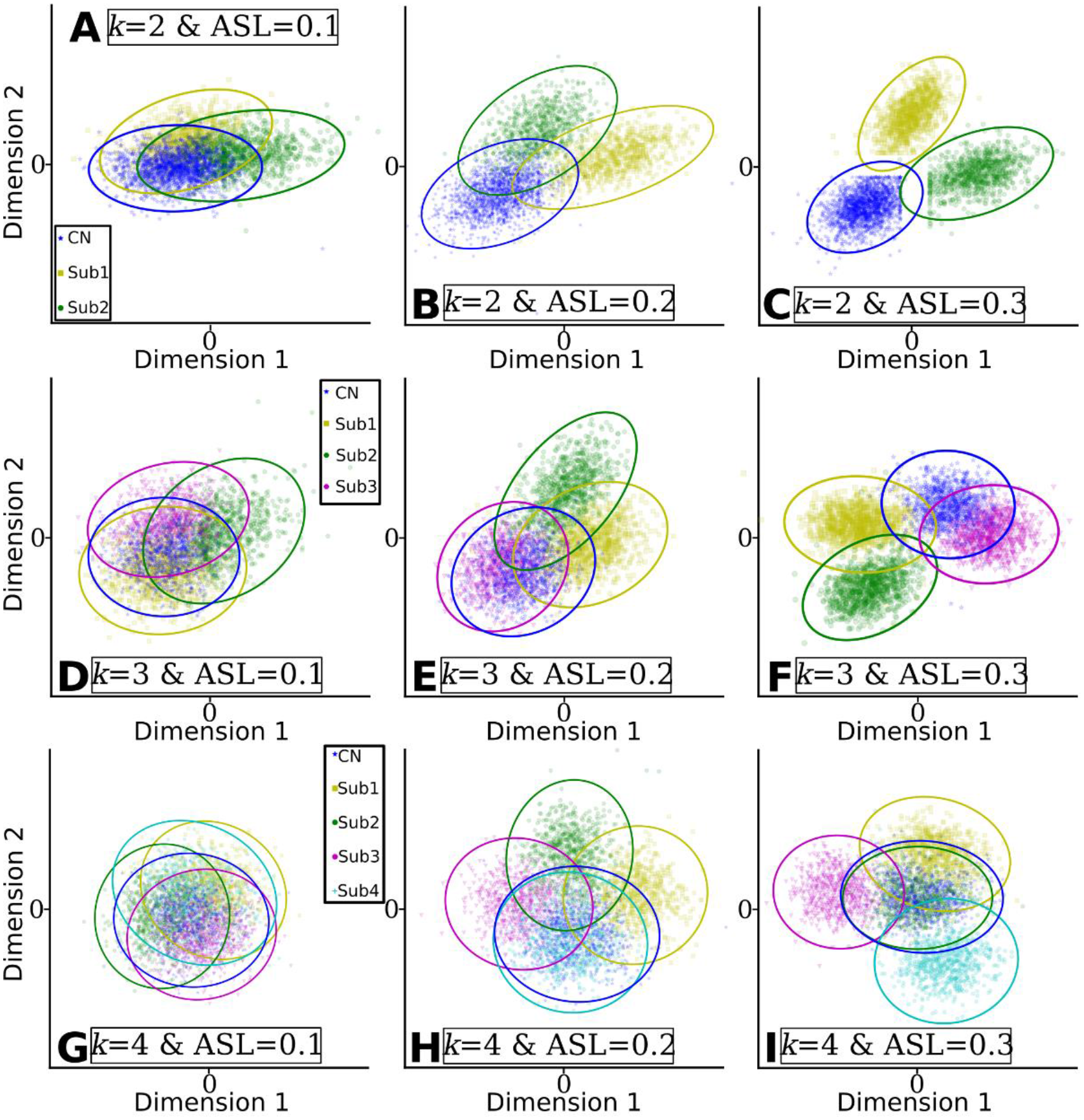
Clusters found by MAGIC become more distinguishable when the clustering conditions are favorable, i.e., higher ASL or lower k. The clusters were projected into the 2D space for visualization. Dimension 1 and Dimension 2 represent the two components projected by multidimensional scaling methods. A) *k*=2 & ASL=0.1; B) *k*=2 & ASL=0.2; C) *k*=2 & ASL=0.3; D) *k*=3 & ASL=0.1; E) *k*=3 & ASL=0.2; F) *k*=3 & ASL=0.3; G) *k*=4 & ASL=0.1; H) *k*=4 & ASL=0.2; I) *k*=4 & ASL=0.3.

**eFigure 5.**
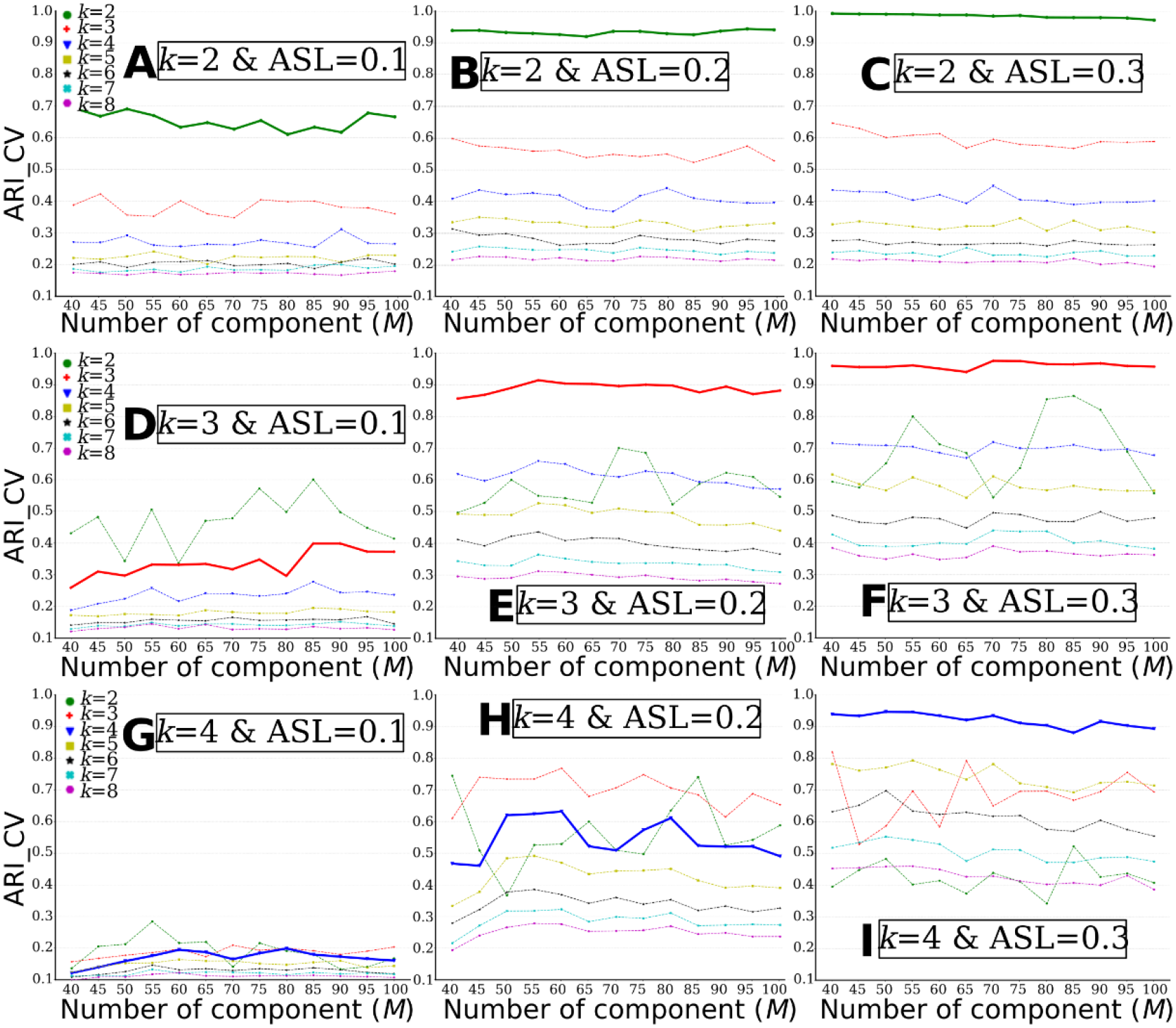
MAGIC finds the ground truth of the number of clusters (*k*) when the clustering conditions are favorable, i.e., higher ASL or lower *k*. The “optimal” k was determined based on the ARI (ARI_CV). A) *k*=2 & ASL=0.1; B) *k*=2 & ASL=0.2; C) *k*=2 & ASL=0.3; D) *k*=3 & ASL=0.1; E) *k*=3 & ASL=0.2; F) *k*=3 & ASL=0.3; G) *k*=4 & ASL=0.1; H) *k*=4 & ASL=0.2; I) *k*=4 & ASL=0.3. The bold lines represent the ground truth of *k* for each experiment.

**eFigure 6.**
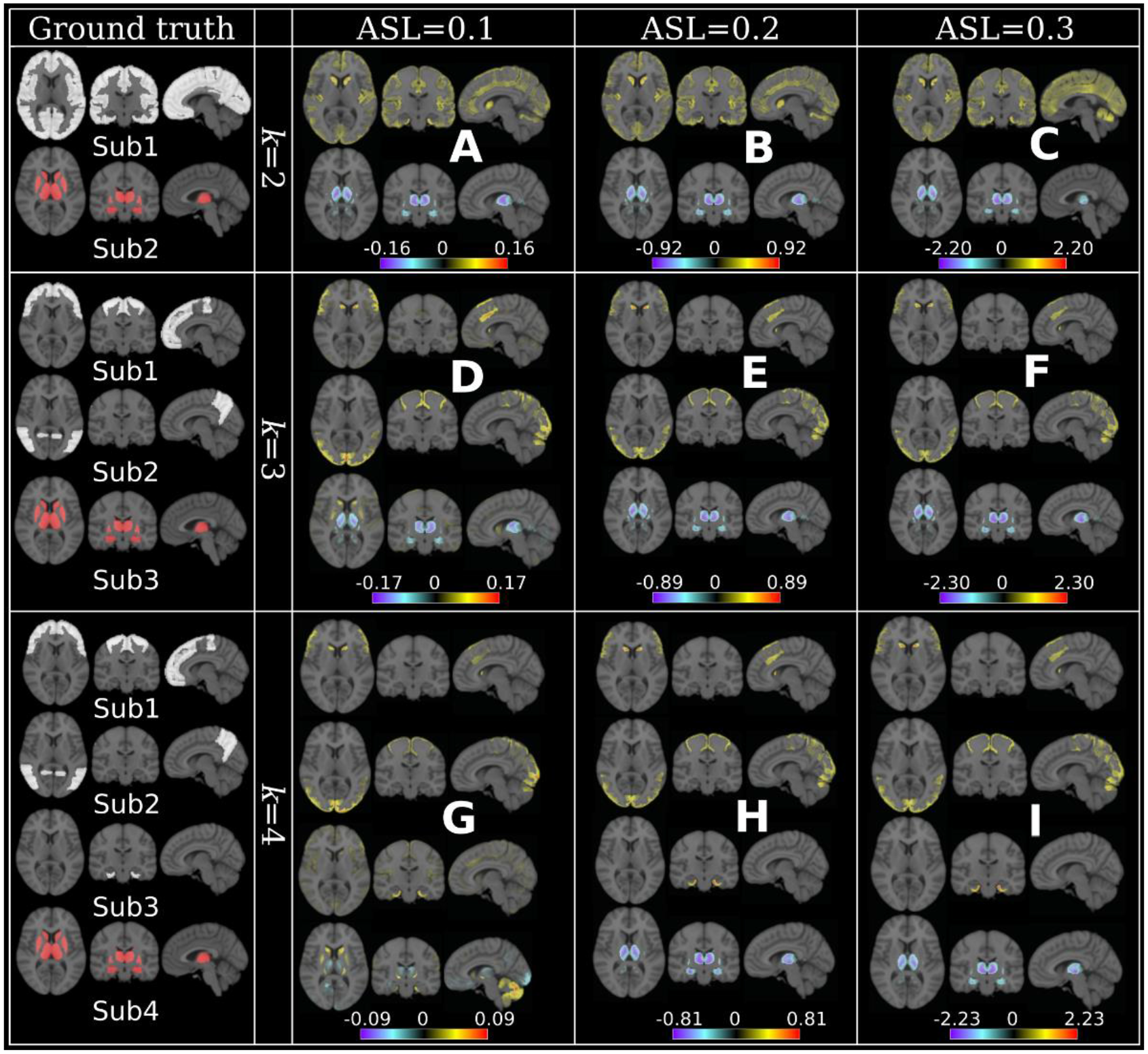
MAGIC finds the ground truth of subtype’s neuroanatomical patterns when the clustering conditions are favorable, i.e., higher ASL or lower *k*. Neuroanatomical patterns are displayed using effect size maps based on voxel-wise group comparisons between CN and subtypes. Positive values denote brain atrophy (CN > Sub), while negative values correspond to larger brain volume in subtypes (CN < Sub). The ground truth of the subtypes pattern is presented with a binary mask (white for positive and red for negative direction) for each *k* in the first column. A) *k*=2 & ASL=0.1; B) *k*=2 & ASL=0.2; C) *k*=2 & ASL=0.3; D) *k*=3 & ASL=0.1; E) *k*=3 & ASL=0.2; F) *k*=3 & ASL=0.3; G) *k*=4 & ASL=0.1; H) *k*=4 & ASL=0.2; I) *k*=4 & ASL=0.3. For reference, Cohen’s *f*2 ≥ 0.02, ≥ 0.15, and ≥ 0.35 represent small, medium, and large effect sizes, respectively.

**eFigure 7.**
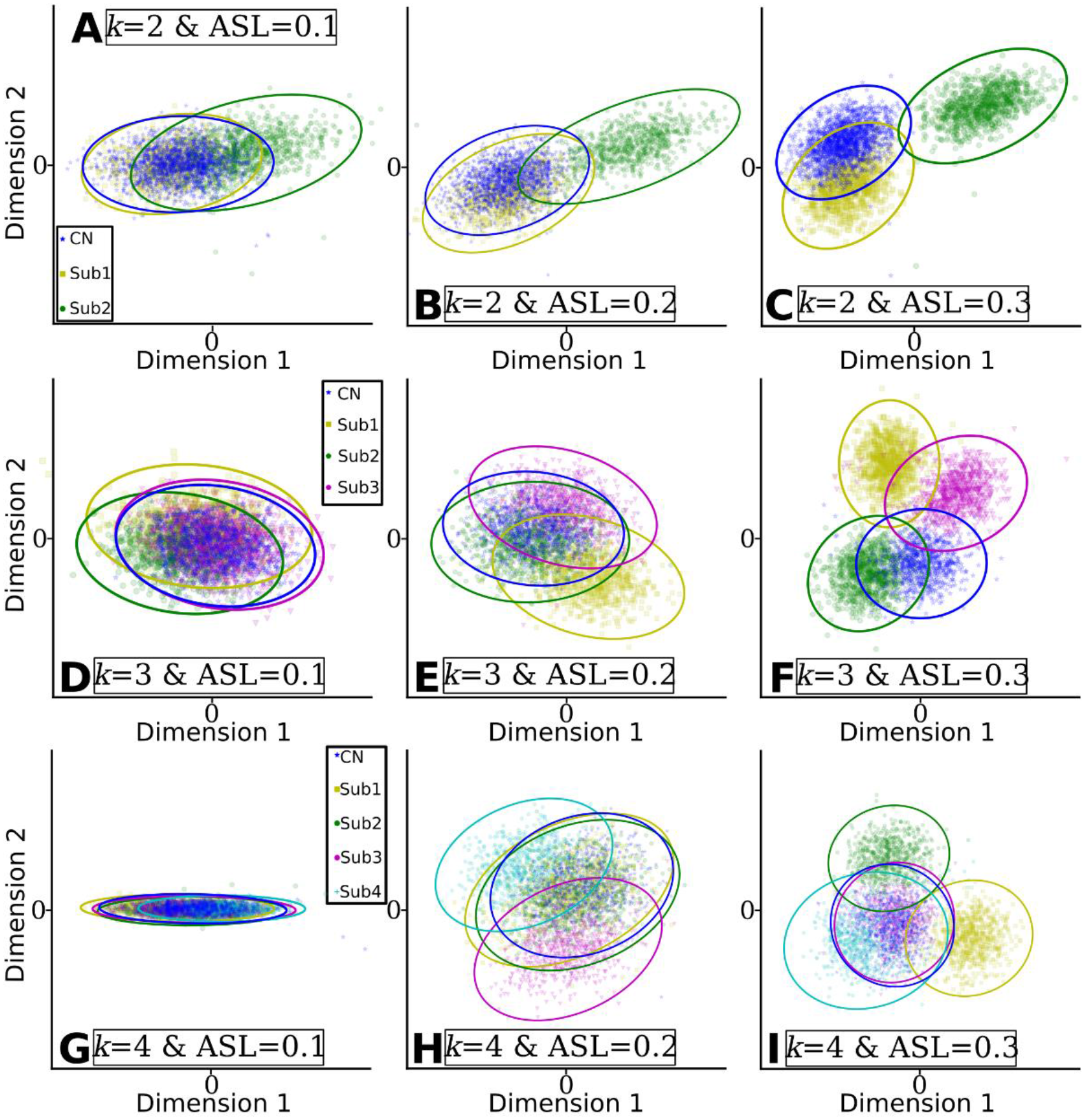
Clusters found by MAGIC become more distinguishable when the clustering conditions are favorable, i.e., higher ASL or lower k. The clusters were projected into the 2D space for visualization. Dimension 1 and Dimension 2 represent the two components projected by multidimensional scaling methods. A) *k*=2 & ASL=0.1; B) *k*=2 & ASL=0.2; C) *k*=2 & ASL=0.3; D) *k*=3 & ASL=0.1; E) *k*=3 & ASL=0.2; F) *k*=3 & ASL=0.3; G) *k*=4 & ASL=0.1; H) *k*=4 & ASL=0.2; I) *k*=4 & ASL=0.3.

**eFigure 8.**
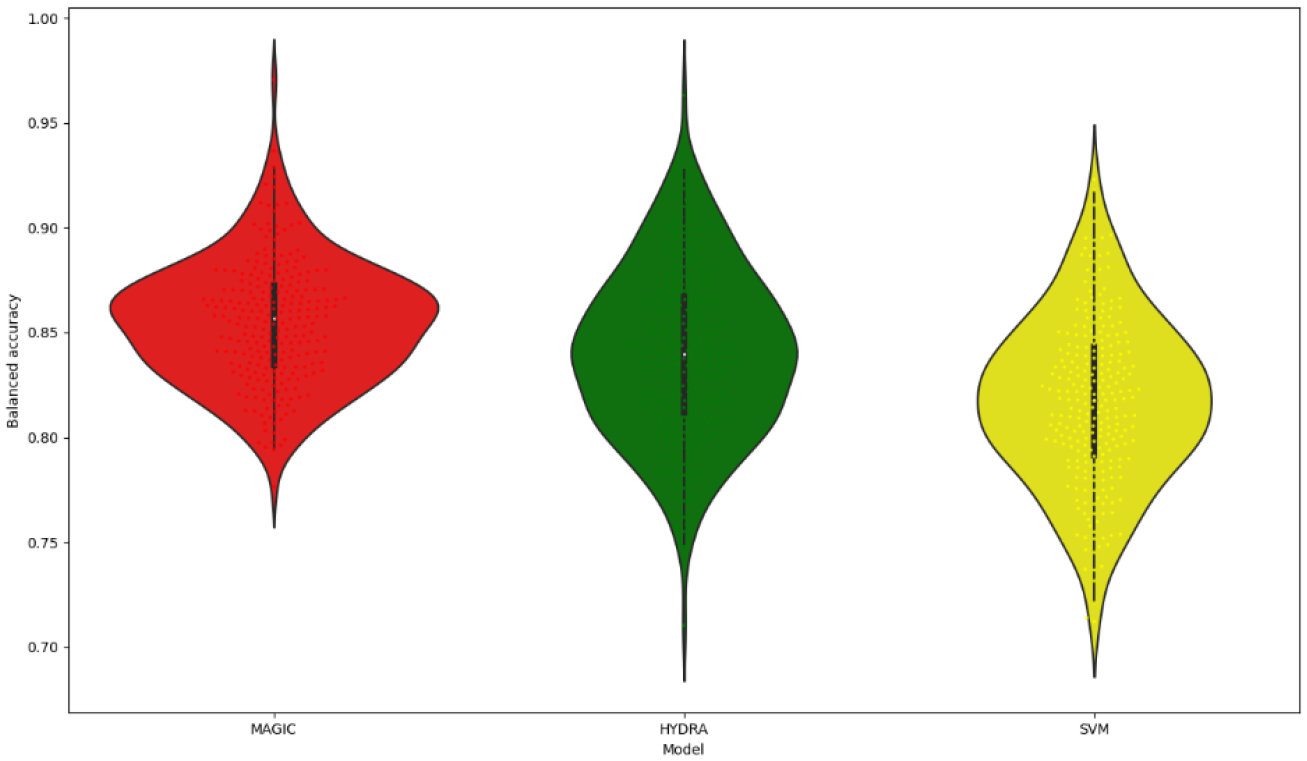
Classification performance for ADNI CN vs AD task across different models, i.e., MAGIC, HYDRA, and a linear SVM. We adopted the same cross-validation (CV) procedure as MAGIC for all models for a fair comparison. Specifically, a non-nested CV with repeated and stratified random splits for 250 repetitions was performed. During each repetition, 80% of the data was for training. We did not adopt a nested CV procedure as in (Samper-González et al., 2018) because this is computationally heavy and technically complex for MAGIC. Therefore, the clustering accuracies reported here are lower than their reproducible baseline performance. For MAGIC, the final polytope (created by 4 SVM hyperplanes) of each repetition was used to classify CN vs AD. MAGIC: 0.85±0.03; HYDRA: 0.84±0.04 and SVM: 0.82±0.04.

**eTable 1.**
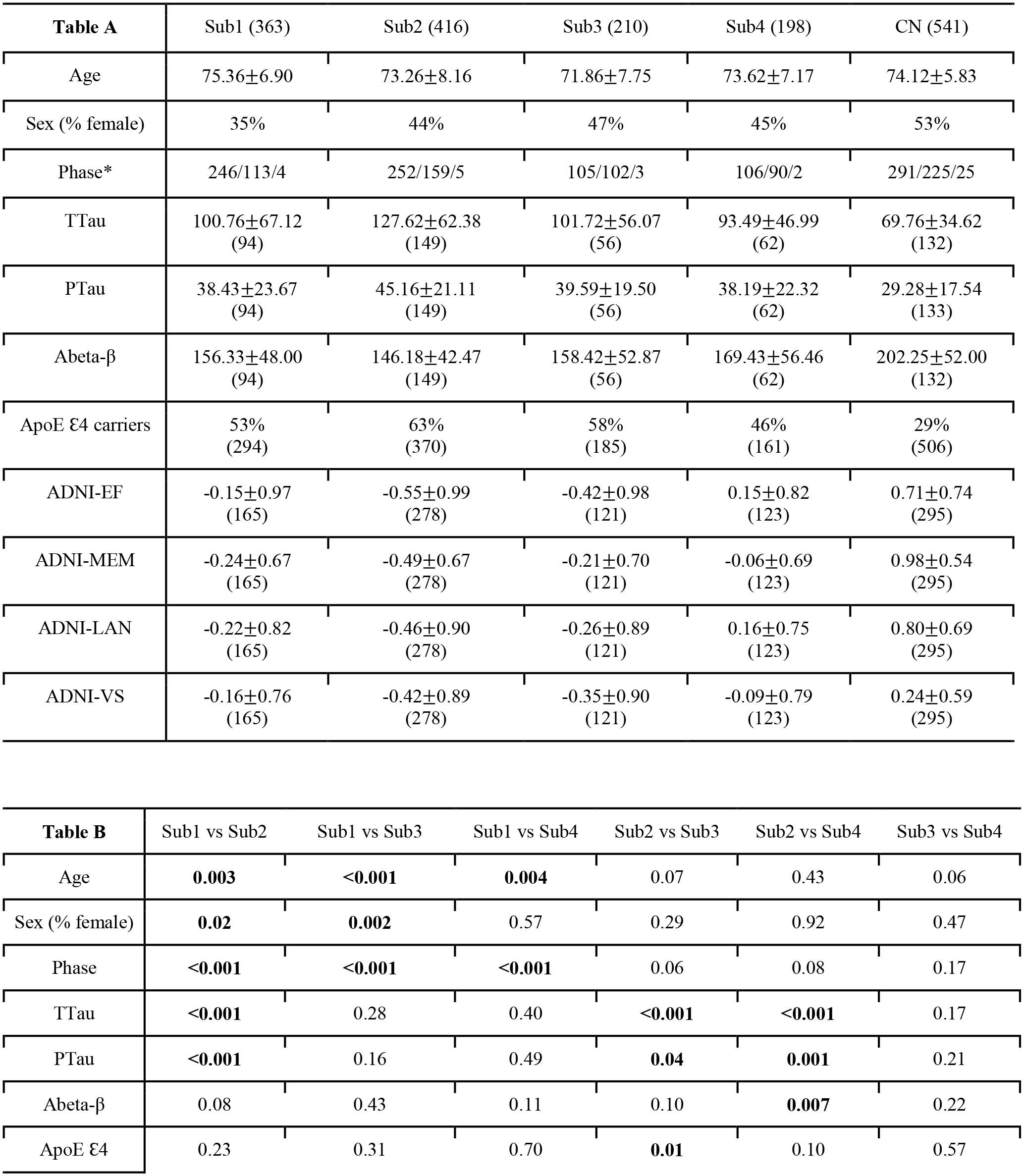
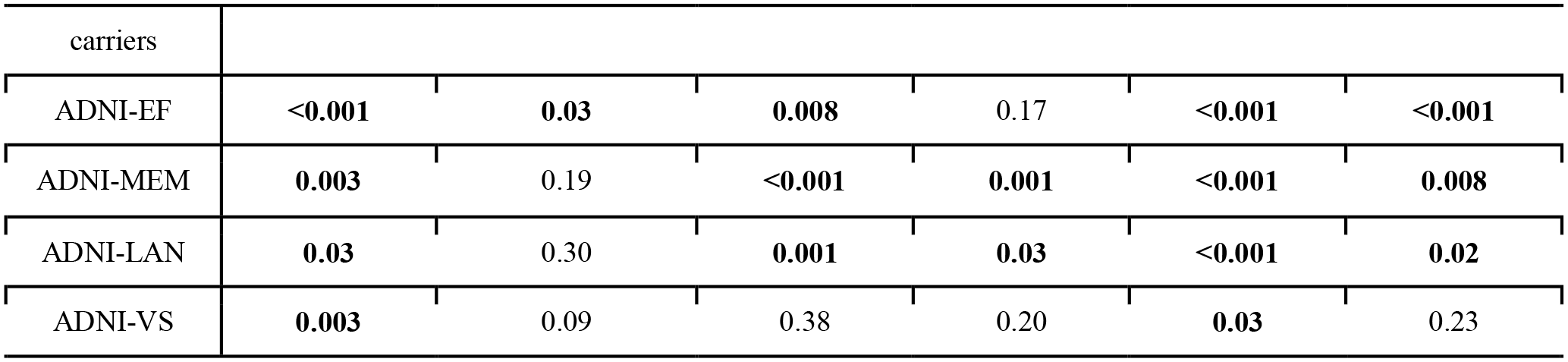
Clinical characteristics of the 4 subtypes found by MAGIC in ADNI. Table A: For a continuous variable, mean and standard deviation are shown; for categorical variables, the percentage is shown. The number of subjects that are available for the data is shown in parentheses. Composite cognitive scores across several domains have been previously validated in ADNI, including a memory composite (ADNI-MEM) (Crane et al., 2012), an executive function composite (ADNI-EF) (Gibbons et al., 2012), a language composite (ADNI-LAN) (Deters et al., 2017), and a visuospatial composite (ADNI-VS) (Choi et al., 2020). For ADNI, data are from three phases, ADNI1, 2, and 3. In Table A, the number was shown for each phase (ADNI1/ADNI2/ADNI3). Table B: We computed the P-value pairwise across the four subtypes. Mann–Whitney–Wilcoxon test was used for continuous variables (e.g., age) and the Chi-Square test of independence for categorical variables (e.g., sex). The significance threshold was 0.05.

**eTable 2.**
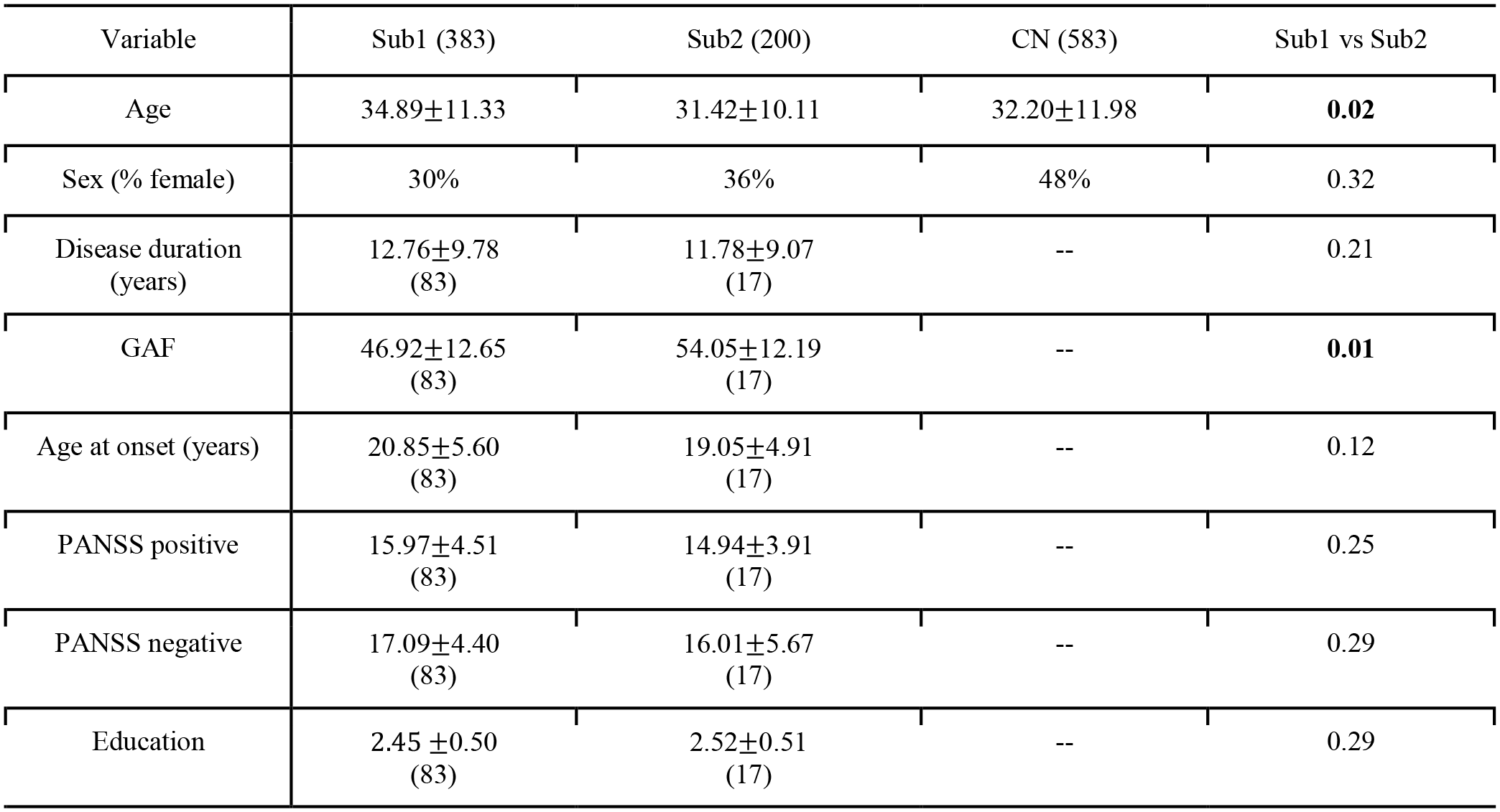
Clinical characteristics of the 2 subtypes found by MAGIC in PHENOM. For a continuous variable, mean and standard deviation are shown; for a categorical variable, the percentage is shown. The number of subjects that are available for the data is shown in parentheses. We computed the P-value pairwise across the two subtypes. Mann–Whitney–Wilcoxon test was used for continuous variables (i.e., age) and ordinal variable (i.e., education), and the Chi-Square test of independence for categorical variables (e.g., sex). The significance threshold was 0.05. -- denotes data not available; PANSS: positive and negative syndrome scale; GAF: global assessment of functioning.

| Variable | Sub1 (383) | Sub2 (200) | CN (583) | Sub1 vs Sub2 |
| --- | --- | --- | --- | --- |
| Age | 34.89±11.33 | 31.42±10.11 | 32.20±11.98 | **0.02** |
| Sex (% female) | 30% | 36% | 48% | 0.32 |
| Disease duration (years) | 12.76±9.78<br>(83) | 11.78±9.07<br>(17) | -- | 0.21 |
| GAF | 46.92±12.65<br>(83) | 54.05±12.19<br>(17) | -- | **0.01** |
| Age at onset (years) | 20.85±5.60<br>(83) | 19.05±4.91<br>(17) | -- | 0.12 |
| PANSS positive | 15.97±4.51<br>(83) | 14.94±3.91<br>(17) | -- | 0.25 |
| PANSS negative | 17.09±4.40<br>(83) | 16.01±5.67<br>(17) | -- | 0.29 |
| Education | 2.45 ±0.50<br>(83) | 2.52±0.51<br>(17) | -- | 0.29 |

## Footnotes

b The term semi-supervised refers to the lack of subtype labels and the use of CN as a reference group to guide the clustering.

c Data used in preparation of this article were obtained from the Alzheimer’s Disease Neuroimaging Initiative (ADNI) database (adni.loni.usc.edu). As such, the investigators within the ADNI contributed to the design and implementation of ADNI and/or provided data but did not participate in analysis or writing of this report. A complete listing of ADNI investigators can be found at: http://adni.loni.usc.edu/wp-content/uploads/how_to_apply/ADNI_Acknowledgement_List.pdf

## References

Abdulkadir, A., Mortamet, B., Vemuri, P., Jack, C.R., Krueger, G., Klöppel, S., Alzheimer’s Disease Neuroimaging Initiative, 2011. Effects of hardware heterogeneity on the performance of SVM Alzheimer’s disease classifier. Neuroimage 58, 785–792. https://doi.org/10.1016/j.neuroimage.2011.06.029

Aisen, P.S., Petersen, R.C., Donohue, M.C., Gamst, A., Raman, R., Thomas, R.G., Walter, S., Trojanowski, J.Q., Shaw, L.M., Beckett, L.A., Jack, C.R., Jr, Jagust, W., Toga, A.W., Saykin, A.J., Morris, J.C., Green, R.C., Weiner, M.W., Alzheimer’s Disease Neuroimaging Initiative, 2010. Clinical Core of the Alzheimer’s Disease Neuroimaging Initiative: progress and plans. Alzheimers. Dement. 6, 239–246.

Altman, N., Krzywinski, M., 2017. Clustering. Nat Methods 14, 545–546. https://doi.org/10.1038/nmeth.4299

Ashburner, J., Friston, K.J., 2000. Voxel-based morphometry--the methods. Neuroimage 11, 805–821. https://doi.org/10.1006/nimg.2000.0582

Ashburner, J., Hutton, C., Frackowiak, R., Johnsrude, I., Price, C., Friston, K., 1998. Identifying global anatomical differences: deformation-based morphometry. Hum Brain Mapp 6, 348–357.

Bashyam, V.M., Erus, G., Doshi, J., Habes, M., Nasrallah, I.M., Truelove-Hill, M., Srinivasan, D., Mamourian, L., Pomponio, R., Fan, Y., Launer, L.J., Masters, C.L., Maruff, P., Zhuo, C., Völzke, H., Johnson, S.C., Fripp, J., Koutsouleris, N., Satterthwaite, T.D., Wolf, D., Gur, R.E., Gur, R.C., Morris, J., Albert, M.S., Grabe, H.J., Resnick, S., Bryan, R.N., Wolk, D.A., Shou, H., Davatzikos, C., on behalf of the ISTAGING Consortium, the P.A. disease C., ADNI, and CARDIA studies, 2020. MRI signatures of brain age and disease over the lifespan based on a deep brain network and 14 468 individuals worldwide. Brain 143, 2312–2324. https://doi.org/10.1093/brain/awaa160

Bassett, D.S., Siebenhühner, F., 2013. Multiscale Network Organization in the Human Brain, in: Multiscale Analysis and Nonlinear Dynamics. John Wiley & Sons, Ltd, pp. 179–204. https://doi.org/10.1002/9783527671632.ch07

Bauermeister, S., Orton, C., Thompson, S., Barker, R.A., Bauermeister, J.R., Ben-Shlomo, Y., Brayne, C., Burn, D., Campbell, A., Calvin, C., Chandran, S., Chaturvedi, N., Chêne, G., Chessell, I.P., Corbett, A., Davis, D.H.J., Denis, M., Dufouil, C., Elliott, P., Fox, N., Hill, D., Hofer, S.M., Hu, M.T., Jindra, C., Kee, F., Kim, C.-H., Kim, C., Kivimaki, M., Koychev, I., Lawson, R.A., Linden, G.J., Lyons, R.A., Mackay, C., Matthews, P.M., McGuiness, B., Middleton, L., Moody, C., Moore, K., Na, D.L., O’Brien, J.T., Ourselin, S., Paranjothy, S., Park, K.-S., Porteous, D.J., Richards, M., Ritchie, C.W., Rohrer, J.D., Rossor, M.N., Rowe, J.B., Scahill, R., Schnier, C., Schott, J.M., Seo, S.W., South, M., Steptoe, M., Tabrizi, S.J., Tales, A., Tillin, T., Timpson, N.J., Toga, A.W., Visser, P.-J., Wade-Martins, R., Wilkinson, T., Williams, J., Wong, A., Gallacher, J.E.J., 2020. The Dementias Platform UK (DPUK) Data Portal. Eur J Epidemiol 35, 601–611. https://doi.org/10.1007/s10654-020-00633-4

Betzel, R.F., Bassett, D.S., 2017. Multi-scale brain networks. NeuroImage, Functional Architecture of the Brain 160, 73–83. https://doi.org/10.1016/j.neuroimage.2016.11.006

Brugger, S.P., Howes, O.D., 2017. Heterogeneity and Homogeneity of Regional Brain Structure in Schizophrenia. JAMA Psychiatry 74, 1104–1111. https://doi.org/10.1001/jamapsychiatry.2017.2663

Chand, G.B., Dwyer, D.B., Erus, G., Sotiras, A., Varol, E., Srinivasan, D., Doshi, J., Pomponio, R., Pigoni, A., Dazzan, P., Kahn, R.S., Schnack, H.G., Zanetti, M.V., Meisenzahl, E., Busatto, G.F., Crespo-Facorro, B., Pantelis, C., Wood, S.J., Zhuo, C., Shinohara, R.T., Shou, H., Fan, Y., Gur, R.C., Gur, R.E., Satterthwaite, T.D., Koutsouleris, N., Wolf, D.H., Davatzikos, C., 2020. Two distinct neuroanatomical subtypes of schizophrenia revealed using machine learning. Brain 143, 1027–1038. https://doi.org/10.1093/brain/awaa025

Chang, C.-C., Lin, C.-J., 2011. LIBSVM: A library for support vector machines. ACM Trans. Intell. Syst. Technol. 2, 1–27. https://doi.org/10.1145/1961189.1961199

Choi, S., Mukherjee, S., Gibbons, L.E., Sanders, R.E., Jones, R.N., Tommet, D., Mez, J., Trittschuh, E.H., Saykin, A., Lamar, M., Rabin, L., Foldi, N.S., Sikkes, S., Jutten, R.J., Grandoit, E., Mac Donald, C., Risacher, S., Groot, C., Ossenkoppele, R., Crane, P.K., 2020. Development and validation of language and visuospatial composite scores in ADNI. Alzheimers Dement (N Y) 6, e12072. https://doi.org/10.1002/trc2.12072

Chu, C., Hsu, A.-L., Chou, K.-H., Bandettini, P.A., Lin, C., Initiative, A.D.N., 2011. Does feature selection improve classification accuracy? Impact of sample size and feature selection on classification using anatomical magnetic resonance images [WWW Document]. NeuroImage. https://doi.org/10.1016/j.neuroimage.2011.11.066

Climescu-Haulica, A., 2007. How to Choose the Number of Clusters: The Cramer Multiplicity Solution, in: Decker, R., Lenz, H.-J. (Eds.), Advances in Data Analysis, Studies in Classification, Data Analysis, and Knowledge Organization. Springer, Berlin, Heidelberg, pp. 15–22. https://doi.org/10.1007/978-3-540-70981-7_2

Cox, M.A.A., Cox, T.F., 2008. Multidimensional Scaling, in: Chen, C., Härdle, W., Unwin, A. (Eds.), Handbook of Data Visualization, Springer Handbooks Comp.Statistics. Springer, Berlin, Heidelberg, pp. 315–347. https://doi.org/10.1007/978-3-540-33037-0_14

Cox, R.W., 1996. AFNI: software for analysis and visualization of functional magnetic resonance neuroimages. Comput. Biomed. Res. 29, 162–173.

Cox, R.W., Chen, G., Glen, D.R., Reynolds, R.C., Taylor, P.A., 2017. fMRI clustering and false-positive rates. Proc Natl Acad Sci U S A 114, E3370–E3371. https://doi.org/10.1073/pnas.1614961114

Crane, P.K., Carle, A., Gibbons, L.E., Insel, P., Mackin, R.S., Gross, A., Jones, R.N., Mukherjee, S., Curtis, S.M., Harvey, D., Weiner, M., Mungas, D., for the Alzheimer’s Disease Neuroimaging Initiative, 2012. Development and assessment of a composite score for memory in the Alzheimer’s Disease Neuroimaging Initiative (ADNI). Brain Imaging and Behavior 6, 502–516. https://doi.org/10.1007/s11682-012-9186-z

Cui, Z., Chen, W., Chen, Y., 2016. Multi-Scale Convolutional Neural Networks for Time Series Classification. ArXiv.

Cuingnet, R., Gerardin, E., Tessieras, J., Auzias, G., Lehéricy, S., Habert, M.-O., Chupin, M., Benali, H., Colliot, O., 2011. Automatic classification of patients with Alzheimer’s disease from structural MRI: A comparison of ten methods using the ADNI database. NeuroImage 56, 766–781. https://doi.org/10.1016/j.neuroimage.2010.06.013

Davatzikos, C., 2019. Machine learning in neuroimaging: Progress and challenges. NeuroImage 197, 652–656. https://doi.org/10.1016/j.neuroimage.2018.10.003

Davatzikos, C., Genc, A., Xu, D., Resnick, S.M., 2001. Voxel-based morphometry using the RAVENS maps: methods and validation using simulated longitudinal atrophy. Neuroimage 14, 1361–1369. https://doi.org/10.1006/nimg.2001.0937

Day, W.H.E., Edelsbrunner, H., 1984. Efficient algorithms for agglomerative hierarchical clustering methods. Journal of Classification 1, 7–24. https://doi.org/10.1007/BF01890115

Deters, K.D., Nho, K., Risacher, S.L., Kim, S., Ramanan, V.K., Crane, P.K., Apostolova, L.G., Saykin, A.J., Alzheimer’s Disease Neuroimaging Initiative, 2017. Genome-wide association study of language performance in Alzheimer’s disease. Brain Lang 172, 22–29. https://doi.org/10.1016/j.bandl.2017.04.008

DeTure, M.A., Dickson, D.W., 2019. The neuropathological diagnosis of Alzheimer’s disease. Molecular Neurodegeneration 14, 32. https://doi.org/10.1186/s13024-019-0333-5

Dong, A., Honnorat, N., Gaonkar, B., Davatzikos, C., 2016a. CHIMERA: Clustering of Heterogeneous Disease Effects via Distribution Matching of Imaging Patterns. IEEE Trans. Med. Imaging 35, 612–621. https://doi.org/10.1109/TMI.2015.2487423

Dong, A., Toledo, J.B., Honnorat, N., Doshi, J., Varol, E., Sotiras, A., Wolk, D., Trojanowski, J.Q., Davatzikos, C., for the Alzheimer’s Disease Neuroimaging Initiative, 2016b. Heterogeneity of neuroanatomical patterns in prodromal Alzheimer’s disease: links to cognition, progression and biomarkers. Brain aww319. https://doi.org/10.1093/brain/aww319

Doshi, J., Erus, G., Ou, Y., Resnick, S.M., Gur, R.C., Gur, R.E., Satterthwaite, T.D., Furth, S., Davatzikos, C., Alzheimer’s Neuroimaging Initiative, 2016. MUSE: MUlti-atlas region Segmentation utilizing Ensembles of registration algorithms and parameters, and locally optimal atlas selection. Neuroimage 127, 186–195. https://doi.org/10.1016/j.neuroimage.2015.11.073

Dubey, R., Zhou, J., Wang, Y., Thompson, P.M., Ye, J., 2014. ANALYSIS OF SAMPLING TECHNIQUES FOR IMBALANCED DATA: AN N=648 ADNI STUDY. Neuroimage 87, 220–241. https://doi.org/10.1016/j.neuroimage.2013.10.005

Dwyer, D.B., Cabral, C., Kambeitz-Ilankovic, L., Sanfelici, R., Kambeitz, J., Calhoun, V., Falkai, P., Pantelis, C., Meisenzahl, E., Koutsouleris, N., 2018. Brain Subtyping Enhances The Neuroanatomical Discrimination of Schizophrenia. Schizophrenia Bulletin 44, 1060–1069. https://doi.org/10.1093/schbul/sby008

Ecker, C., Rocha-Rego, V., Johnston, P., Mourao-Miranda, J., Marquand, A., Daly, E.M., Brammer, M.J., Murphy, C., Murphy, D.G., MRC AIMS Consortium, 2010. Investigating the predictive value of whole-brain structural MR scans in autism: a pattern classification approach. Neuroimage 49, 44–56. https://doi.org/10.1016/j.neuroimage.2009.08.024

Ezzati, A., Ezzati, A., Zammit, A.R., Habeck, C., Hall, C.B., Lipton, R.B., 2020. Detecting biological heterogeneity patterns in ADNI amnestic mild cognitive impairment based on volumetric MRI. Brain Imaging and Behavior 14, 1792–1804. https://doi.org/10.1007/s11682-019-00115-6

Filipovych, R., Resnick, S.M., Davatzikos, C., 2012. JointMMCC: Joint Maximum-Margin Classification and Clustering of Imaging Data. IEEE Trans. Med. Imaging 31, 1124–1140. https://doi.org/10.1109/TMI.2012.2186977

Franke, K., Ziegler, G., Klöppel, S., Gaser, C., Alzheimer’s Disease Neuroimaging Initiative, 2010. Estimating the age of healthy subjects from T1-weighted MRI scans using kernel methods: exploring the influence of various parameters. Neuroimage 50, 883–892. https://doi.org/10.1016/j.neuroimage.2010.01.005

Friston, K.J., Holmes, A.P., Worsley, K.J., Poline, J.-P., Frith, C.D., Frackowiak, R.S.J., 1994. Statistical parametric maps in functional imaging: A general linear approach. Human Brain Mapping 2, 189–210. https://doi.org/10.1002/hbm.460020402

Fu, W., Perry, P.O., 2020. Estimating the Number of Clusters Using Cross-Validation. Journal of Computational and Graphical Statistics 29, 162–173. https://doi.org/10.1080/10618600.2019.1647846

Gaonkar, B., Davatzikos, C., 2013. Analytic estimation of statistical significance maps for support vector machine based multi-variate image analysis and classification. NeuroImage 78, 270–283. https://doi.org/10.1016/j.neuroimage.2013.03.066

Gibbons, L.E., Carle, A.C., Mackin, R.S., Harvey, D., Mukherjee, S., Insel, P., Curtis, S.M., Mungas, D., Crane, P.K., Alzheimer’s Disease Neuroimaging Initiative, 2012. A composite score for executive functioning, validated in Alzheimer’s Disease Neuroimaging Initiative (ADNI) participants with baseline mild cognitive impairment. Brain Imaging Behav 6, 517–527. https://doi.org/10.1007/s11682-012-9176-1

Habes, M., Janowitz, D., Erus, G., Toledo, J.B., Resnick, S.M., Doshi, J., Van der Auwera, S., Wittfeld, K., Hegenscheid, K., Hosten, N., Biffar, R., Homuth, G., Völzke, H., Grabe, H.J., Hoffmann, W., Davatzikos, C., 2016. Advanced brain aging: relationship with epidemiologic and genetic risk factors, and overlap with Alzheimer disease atrophy patterns. Transl Psychiatry 6, e775–e775. https://doi.org/10.1038/tp.2016.39

Hanyu, H., Sakurai, H., Iwamoto, T., Takasaki, M., Shindo, H., Abe, K., 1998. Diffusion-weighted MR imaging of the hippocampus and temporal white matter in Alzheimer’s disease. J. Neurol. Sci. 156, 195–200.

Hartigan, J.A., Wong, M.A., 1979. Algorithm AS 136: A K-Means Clustering Algorithm. Journal of the Royal Statistical Society. Series C (Applied Statistics) 28, 100–108. https://doi.org/10.2307/2346830

Honnorat, N., Dong, A., Meisenzahl-Lechner, E., Koutsouleris, N., Davatzikos, C., 2019. Neuroanatomical heterogeneity of schizophrenia revealed by semi-supervised machine learning methods. Schizophrenia Research 214, 43–50. https://doi.org/10.1016/j.schres.2017.12.008

Hu, K., Wang, Y., Chen, K., Hou, L., Zhang, X., 2016. Multi-scale features extraction from baseline structure MRI for MCI patient classification and AD early diagnosis. Neurocomputing 175, 132–145. https://doi.org/10.1016/j.neucom.2015.10.043

Insel, T.R., Cuthbert, B.N., 2015. Brain disorders? Precisely. Science 348, 499–500. https://doi.org/10.1126/science.aab2358

Jack, C.R., Bennett, D.A., Blennow, K., Carrillo, M.C., Feldman, H.H., Frisoni, G.B., Hampel, H., Jagust, W.J., Johnson, K.A., Knopman, D.S., Petersen, R.C., Scheltens, P., Sperling, R.A., Dubois, B., 2016. A/T/N: An unbiased descriptive classification scheme for Alzheimer disease biomarkers. Neurology 87, 539– 547. https://doi.org/10.1212/WNL.0000000000002923

Jeon, S., Kang, J.M., Seo, S., Jeong, H.J., Funck, T., Lee, S.-Y., Park, K.H., Lee, Y.-B., Yeon, B.K., Ido, T., Okamura, N., Evans, A.C., Na, D.L., Noh, Y., 2019. Topographical Heterogeneity of Alzheimer’s Disease Based on MR Imaging, Tau PET, and Amyloid PET. Front. Aging Neurosci. 11, 211. https://doi.org/10.3389/fnagi.2019.00211

Jung, N.-Y., Seo, S.W., Yoo, H., Yang, J.-J., Park, S., Kim, Y.J., Lee, J., Lee, J.S., Jang, Y.K., Lee, J.M., Kim, S.T., Kim, S., Kim, E.-J., Na, D.L., Kim, H.J., 2016. Classifying anatomical subtypes of subjective memory impairment. Neurobiology of Aging 48, 53–60. https://doi.org/10.1016/j.neurobiolaging.2016.08.010

Kamnitsas, K., Ledig, C., Newcombe, V., Simpson, J., Kane, A.D., Menon, D., Rueckert, D., Glocker, B., 2017. Efficient multi-scale 3D CNN with fully connected CRF for accurate brain lesion segmentation. Medical Image Anal. https://doi.org/10.1016/j.media.2016.10.004

Koutsouleris, N., Meisenzahl, E.M., Borgwardt, S., Riecher-Rössler, A., Frodl, T., Kambeitz, J., Köhler, Y., Falkai, P., Möller, H.-J., Reiser, M., Davatzikos, C., 2015. Individualized differential diagnosis of schizophrenia and mood disorders using neuroanatomical biomarkers. Brain 138, 2059–2073. https://doi.org/10.1093/brain/awv111

Lao, Z., Shen, D., Xue, Z., Karacali, B., Resnick, S.M., Davatzikos, C., 2004. Morphological classification of brains via high-dimensional shape transformations and machine learning methods. NeuroImage 21, 46– 57. https://doi.org/10.1016/j.neuroimage.2003.09.027

Lee, D.D., Seung, H.S., 2001. Algorithms for Non-negative Matrix Factorization 7.

Lubeiro, A., Rueda, C., Hernández, J.A., Sanz, J., Sarramea, F., Molina, V., 2016. Identification of two clusters within schizophrenia with different structural, functional and clinical characteristics. Progress in Neuro-Psychopharmacology and Biological Psychiatry 64, 79–86. https://doi.org/10.1016/j.pnpbp.2015.06.015

McLachlan, G.J., Basford, K.E., 1988. Mixture Models: Inference And Applications To Clustering 1.

Miller, K.L., Alfaro-Almagro, F., Bangerter, N.K., Thomas, D.L., Yacoub, E., Xu, J., Bartsch, A.J., Jbabdi, S., Sotiropoulos, S.N., Andersson, J.L., Griffanti, L., Douaud, G., Okell, T.W., Weale, P., Dragonu, I., Garratt, S., Hudson, S., Collins, R., Jenkinson, M., Matthews, P.M., Smith, S.M., 2016. Multimodal population brain imaging in the UK Biobank prospective epidemiological study. Nat Neurosci 19, 1523–1536. https://doi.org/10.1038/nn.4393

Miotto, R., Wang, F., Wang, S., Jiang, X., Dudley, J.T., 2018. Deep learning for healthcare: review, opportunities and challenges. Briefings in Bioinformatics 19, 1236–1246. https://doi.org/10.1093/bib/bbx044

Mirkin, B., 2011. Choosing the number of clusters. WIREs Data Mining Knowl Discov 1, 252–260. https://doi.org/10.1002/widm.15

Müller, M.J., Greverus, D., Dellani, P.R., Weibrich, C., Wille, P.R., Scheurich, A., Stoeter, P., Fellgiebel, A., 2005. Functional implications of hippocampal volume and diffusivity in mild cognitive impairment. Neuroimage 28, 1033–1042.

Murray, M.E., Graff-Radford, N.R., Ross, O.A., Petersen, R.C., Duara, R., Dickson, D.W., 2011. Neuropathologically defined subtypes of Alzheimer’s disease with distinct clinical characteristics: a retrospective study. The Lancet Neurology 10, 785–796. https://doi.org/10.1016/S1474-4422(11)70156-9

Nadeau, C., Bengio, Y., 2003. 46-Inference for the Generalization Error. Machine Learning 52, 239–281. https://doi.org/10.1023/A:1024068626366

Nettiksimmons, J., DeCarli, C., Landau, S., Beckett, L., 2014. Biological heterogeneity in ADNI amnestic mild cognitive impairment. Alzheimer’s & Dementia 10, 511–521.e1. https://doi.org/10.1016/j.jalz.2013.09.003

Ng, A.Y., Jordan, M.I., Weiss, Y., 2001. On Spectral Clustering: Analysis and an algorithm, in: Advances in Neural Information Processing Systems. MIT Press, pp. 849–856.

Noh, Y., Jeon, S., Lee, J.M., Seo, S.W., Kim, G.H., Cho, H., Ye, B.S., Yoon, C.W., Kim, H.J., Chin, J., Park, K.H., Heilman, K.M., Na, D.L., 2014. Anatomical heterogeneity of Alzheimer disease: Based on cortical thickness on MRIs. Neurology 83, 1936–1944. https://doi.org/10.1212/WNL.0000000000001003

Okada, N, Okada, N., Fukunaga, M., Yamashita, F., Koshiyama, D., Yamamori, H., Ohi, K., Yasuda, Y., Fujimoto, M., Watanabe, Y., Yahata, N., Nemoto, K., Hibar, D.P., van Erp, T.G.M., Fujino, H., Isobe, M., Isomura, S., Natsubori, T., Narita, H., Hashimoto, N., Miyata, J., Koike, S., Takahashi, T., Yamasue, H., Matsuo, K., Onitsuka, T., Iidaka, T., Kawasaki, Y., Yoshimura, R., Watanabe, Y., Suzuki, M., Turner, J.A., Takeda, M., Thompson, P.M., Ozaki, N., Kasai, K., Hashimoto, R., 2016. Abnormal asymmetries in subcortical brain volume in schizophrenia. Mol Psychiatry 21, 1460–1466. https://doi.org/10.1038/mp.2015.209

Ota, K., for the Alzheimer’s Disease Neuroimaging Initiative, Ota, K., Oishi, N., Ito, K., Fukuyama, H., 2016. Prediction of Alzheimer’s Disease in Amnestic Mild Cognitive Impairment Subtypes: Stratification Based on Imaging Biomarkers. JAD 52, 1385–1401. https://doi.org/10.3233/JAD-160145

Ou, Y., Sotiras, A., Paragios, N., Davatzikos, C., 2011. DRAMMS: Deformable Registration via Attribute Matching and Mutual-Saliency Weighting. Med Image Anal 15, 622–639. https://doi.org/10.1016/j.media.2010.07.002

Pan, Y., Pu, W., Chen, X., Huang, X., Cai, Y., Tao, H., Xue, Z., Mackinley, M., Limongi, R., Liu, Z., Palaniyappan, L., 2020. Morphological Profiling of Schizophrenia: Cluster Analysis of MRI-Based Cortical Thickness Data. Schizophrenia Bulletin 46, 623–632. https://doi.org/10.1093/schbul/sbz112

Park, J.-Y., Park, J.-Y., Na, H.K., Kim, S., Kim, H., Kim, H.J., Seo, S.W., Na, D.L., Han, C.E., Seong, J.-K., 2017. Robust Identification of Alzheimer’s Disease subtypes based on cortical atrophy patterns. Sci Rep 7, 43270. https://doi.org/10.1038/srep43270

Perl, D.P., 2010. Neuropathology of Alzheimer’s Disease. Mt Sinai J Med 77, 32–42. https://doi.org/10.1002/msj.20157

Petersen, R.C., Aisen, P.S., Beckett, L.A., Donohue, M.C., Gamst, A.C., Harvey, D.J., Jack, C.R., Jr, Jagust, W.J., Shaw, L.M., Toga, A.W., Trojanowski, J.Q., Weiner, M.W., 2010. Alzheimer’s Disease Neuroimaging Initiative (ADNI): clinical characterization. Neurology 74, 201–209.

Planchuelo-Gómez, Á., Lubeiro, A., Núñez-Novo, P., Gomez-Pilar, J., de Luis-García, R., del Valle, P., Martín-Santiago, Ó., Pérez-Escudero, A., Molina, V., 2020. Identificacion of MRI-based psychosis subtypes: Replication and refinement. Progress in Neuro-Psychopharmacology and Biological Psychiatry 100, 109907. https://doi.org/10.1016/j.pnpbp.2020.109907

Pomponio, R., Erus, G., Habes, M., Doshi, J., Srinivasan, D., Mamourian, E., Bashyam, V., Fan, Y., Launer, L.J., Masters, C.L., Maruff, P., Zhuo, C., Nasrallah, I.M., Völzke, H., Johnson, S.C., Fripp, J., Koutsouleris, N., Satterthwaite, T.D., Wolf, D.H., Gur, Raquel, Gur, Ruben, Morris, J., Albert, M.S., Grabe, H.J., Resnick, S.M., Bryan, R.N., Wolk, D.A., Shinohara, R.T., Shou, H., Davatzikos, C., 2019. Harmonization of large multi-site imaging datasets: Application to 10,232 MRIs for the analysis of imaging patterns of structural brain change throughout the lifespan (preprint). Bioinformatics. https://doi.org/10.1101/784363

Poulakis, K., Ferreira, D., Pereira, J.B., Smedby, Ö., Vemuri, P., Westman, E., 2020. Fully bayesian longitudinal unsupervised learning for the assessment and visualization of AD heterogeneity and progression 26.

Poulakis, K., Pereira, J.B., Mecocci, P., Vellas, B., Tsolaki, M., Kłoszewska, I., Soininen, H., Lovestone, S., Simmons, A., Wahlund, L.-O., Westman, E., 2018. Heterogeneous patterns of brain atrophy in Alzheimer’s disease. Neurobiology of Aging 65, 98–108. https://doi.org/10.1016/j.neurobiolaging.2018.01.009

Rabinovici, G.D., Carrillo, M.C., Forman, M., DeSanti, S., Miller, D.S., Kozauer, N., Petersen, R.C., Randolph, C., Knopman, D.S., Smith, E.E., Isaac, M., Mattsson, N., Bain, L.J., Hendrix, J.A., Sims, J.R., 2016. Multiple comorbid neuropathologies in the setting of Alzheimer’s disease neuropathology and implications for drug development. Alzheimers Dement (N Y) 3, 83–91. https://doi.org/10.1016/j.trci.2016.09.002

Rathore, S., Habes, M., Iftikhar, M.A., Shacklett, A., Davatzikos, C., 2017. A review on neuroimaging-based classification studies and associated feature extraction methods for Alzheimer’s disease and its prodromal stages. Neuroimage 155, 530–548.

Rozycki, M., Satterthwaite, T.D., Koutsouleris, N., Erus, G., Doshi, J., Wolf, D.H., Fan, Y., Gur, R.E., Gur, R.C., Meisenzahl, E.M., Zhuo, C., Yin, H., Yan, H., Yue, W., Zhang, D., Davatzikos, C., 2018. Multisite Machine Learning Analysis Provides a Robust Structural Imaging Signature of Schizophrenia Detectable Across Diverse Patient Populations and Within Individuals. Schizophrenia Bulletin 44, 1035–1044. https://doi.org/10.1093/schbul/sbx137

Samper-González, J., Burgos, N., Bottani, S., Fontanella, S., Lu, P., Marcoux, A., Routier, A., Guillon, J., Bacci, M., Wen, J., Bertrand, A., Bertin, H., Habert, M.-O., Durrleman, S., Evgeniou, T., Colliot, O., 2018. Reproducible evaluation of classification methods in Alzheimer’s disease: Framework and application to MRI and PET data. NeuroImage 183, 504–521. https://doi.org/10.1016/j.neuroimage.2018.08.042

Satterthwaite, T.D., Wolf, D.H., Loughead, J., Ruparel, K., Valdez, J.N., Siegel, S.J., Kohler, C.G., Gur, R.E., Gur, R.C., 2010. Association of enhanced limbic response to threat with decreased cortical facial recognition memory response in schizophrenia. Am J Psychiatry 167, 418–426. https://doi.org/10.1176/appi.ajp.2009.09060808

Schirner, M., McIntosh, A.R., Jirsa, V., Deco, G., Ritter, P., 2018. Inferring multi-scale neural mechanisms with brain network modelling. eLife 7, e28927. https://doi.org/10.7554/eLife.28927

Schnack, H.G., Nieuwenhuis, M., van Haren, N.E.M., Abramovic, L., Scheewe, T.W., Brouwer, R.M., Hulshoff Pol, H.E., Kahn, R.S., 2014. Can structural MRI aid in clinical classification? A machine learning study in two independent samples of patients with schizophrenia, bipolar disorder and healthy subjects. Neuroimage 84, 299–306. https://doi.org/10.1016/j.neuroimage.2013.08.053

Schulz, M.-A., Chapman-Rounds, M., Verma, M., Bzdok, D., Georgatzis, K., 2020a. Inferring disease subtypes from clusters in explanation space. Sci Rep 10, 12900. https://doi.org/10.1038/s41598-020-68858-7

Schulz, M.-A., Yeo, B.T.T., Vogelstein, J.T., Mourao-Miranada, J., Kather, J.N., Kording, K., Richards, B., Bzdok, D., 2020b. Different scaling of linear models and deep learning in UKBiobank brain images versus machine-learning datasets. Nat Commun 11, 4238. https://doi.org/10.1038/s41467-020-18037-z

Selya, A.S., Rose, J.S., Dierker, L.C., Hedeker, D., Mermelstein, R.J., 2012. A Practical Guide to Calculating Cohen’s f2, a Measure of Local Effect Size, from PROC MIXED. Front Psychol 3. https://doi.org/10.3389/fpsyg.2012.00111

Sotiras, A., Resnick, S.M., Davatzikos, C., 2015. Finding imaging patterns of structural covariance via Non-Negative Matrix Factorization. NeuroImage 108, 1–16. https://doi.org/10.1016/j.neuroimage.2014.11.045

Starck, J.-L., Murtagh, F., Bijaoui, A., 1998. Image Processing and Data Analysis -The Multiscale Approach. https://doi.org/10.1017/CBO9780511564352

Sugihara, G., Oishi, N., Son, S., Kubota, M., Takahashi, H., Murai, T., 2016. Distinct Patterns of Cerebral Cortical Thinning in Schizophrenia: A Neuroimaging Data-Driven Approach. SCHBUL sbw176. https://doi.org/10.1093/schbul/sbw176

Ten Kate, M., Dicks, E., Visser, P.J., van der Flier, W.M., Teunissen, C.E., Barkhof, F., Scheltens, P., Tijms, B.M., Alzheimer’s Disease Neuroimaging Initiative, 2018. Atrophy subtypes in prodromal Alzheimer’s disease are associated with cognitive decline. Brain 141, 3443–3456. https://doi.org/10.1093/brain/awy264

Tustison, N.J., Avants, B.B., Cook, P.A., Zheng, Y., Egan, A., Yushkevich, P.A., Gee, J.C., 2010. N4ITK: improved N3 bias correction. IEEE Trans. Med. Imaging 29, 1310–1320.

van Erp, T.G.M., Hibar, D.P., Rasmussen, J.M., Glahn, D.C., Pearlson, G.D., Andreassen, O.A., Agartz, I., Westlye, L.T., Haukvik, U.K., Dale, A.M., Melle, I., Hartberg, C.B., Gruber, O., Kraemer, B., Zilles, D., Donohoe, G., Kelly, S., McDonald, C., Morris, D.W., Cannon, D.M., Corvin, A., Machielsen, M.W.J., Koenders, L., de Haan, L., Veltman, D.J., Satterthwaite, T.D., Wolf, D.H., Gur, R.C., Gur, R.E., Potkin, S.G., Mathalon, D.H., Mueller, B.A., Preda, A., Macciardi, F., Ehrlich, S., Walton, E., Hass, J., Calhoun, V.D., Bockholt, H.J., Sponheim, S.R., Shoemaker, J.M., van Haren, N.E.M., Pol, H.E.H., Ophoff, R.A., Kahn, R.S., Roiz-Santiañez, R., Crespo-Facorro, B., Wang, L., Alpert, K.I., Jönsson, E.G., Dimitrova, R., Bois, C., Whalley, H.C., McIntosh, A.M., Lawrie, S.M., Hashimoto, R., Thompson, P.M., Turner, J.A., 2016. Subcortical brain volume abnormalities in 2028 individuals with schizophrenia and 2540 healthy controls via the ENIGMA consortium. Mol Psychiatry 21, 547–553. https://doi.org/10.1038/mp.2015.63

Varghese, T., Sheelakumari, R., James, J.S., Mathuranath, P., 2013. A review of neuroimaging biomarkers of Alzheimer’s disease. Neurol Asia 18, 239–248.

Varol, E., Sotiras, A., Davatzikos, C., 2018. MIDAS: Regionally linear multivariate discriminative statistical mapping. NeuroImage 174, 111–126. https://doi.org/10.1016/j.neuroimage.2018.02.060

Varol, E., Sotiras, A., Davatzikos, C., 2017. HYDRA: Revealing heterogeneity of imaging and genetic patterns through a multiple max-margin discriminative analysis framework. NeuroImage 145, 346–364. https://doi.org/10.1016/j.neuroimage.2016.02.041

Wen, J., Thibeau-Sutre, E., Diaz-Melo, M., Samper-González, J., Routier, A., Bottani, S., Dormont, D., Durrleman, S., Burgos, N., Colliot, O., 2020a. Convolutional neural networks for classification of Alzheimer’s disease: Overview and reproducible evaluation. Medical Image Analysis 63, 101694. https://doi.org/10.1016/j.media.2020.101694

Wen, J., Varol, E., Chand, G., Sotiras, A., Davatzikos, C., 2020b. MAGIC: Multi-scale Heterogeneity Analysis and Clustering for Brain Diseases, in: Martel, A.L., Abolmaesumi, P., Stoyanov, D., Mateus, D., Zuluaga, M.A., Zhou, S.K., Racoceanu, D., Joskowicz, L. (Eds.), Medical Image Computing and Computer Assisted Intervention – MICCAI 2020, Lecture Notes in Computer Science. Springer International Publishing, Cham, pp. 678–687. https://doi.org/10.1007/978-3-030-59728-3_66

Whitwell, J.L., Petersen, R.C., Negash, S., Weigand, S.D., Kantarci, K., Ivnik, R.J., Knopman, D.S., Boeve, B.F., Smith, G.E., Jack, C.R., 2007. Patterns of Atrophy differ among Specific Subtypes of Mild Cognitive Impairment. Arch Neurol 64, 1130–1138. https://doi.org/10.1001/archneur.64.8.1130

Wolf, D.H., Satterthwaite, T.D., Kantrowitz, J.J., Katchmar, N., Vandekar, L., Elliott, M.A., Ruparel, K., 2014. Amotivation in Schizophrenia: Integrated Assessment With Behavioral, Clinical, and Imaging Measures. Schizophr Bull 40, 1328–1337. https://doi.org/10.1093/schbul/sbu026

Wood, S.J., Velakoulis, D., Smith, D.J., Bond, D., Stuart, G.W., McGorry, P.D., Brewer, W.J., Bridle, N., Eritaia, J., Desmond, P., Singh, B., Copolov, D., Pantelis, C., 2001. A longitudinal study of hippocampal volume in first episode psychosis and chronic schizophrenia. Schizophr Res 52, 37–46. https://doi.org/10.1016/s0920-9964(01)00175-x

Yang, Z., Nasrallah, I.M., Shou, H., Wen, J., Doshi, J., Habes, M., Erus, G., Abdulkadir, A., Resnick, S.M., Wolk, D., Davatzikos, C., 2021. Disentangling brain heterogeneity via semi-supervised deep-learning and MRI: dimensional representations of Alzheimer’s Disease. arXiv:2102.12582 [cs, eess, q-bio].

Yang, Z., Wen, J., Davatzikos, C., 2020. Smile-GANs: Semi-supervised clustering via GANs for dissecting brain disease heterogeneity from medical images. arXiv:2006.15255 [cs, eess, q-bio, stat].

Young, A.L., The Alzheimer’s Disease Neuroimaging Initiative (ADNI), Young, A.L., Marinescu, R.V., Oxtoby, N.P., Bocchetta, M., Yong, K., Firth, N.C., Cash, D.M., Thomas, D.L., Dick, K.M., Cardoso, J., van Swieten, J., Borroni, B., Galimberti, D., Masellis, M., Tartaglia, M.C., Rowe, J.B., Graff, C., Tagliavini, F., Frisoni, G.B., Laforce, R., Finger, E., de Mendonça, A., Sorbi, S., Warren, J.D., Crutch, S., Fox, N.C., Ourselin, S., Schott, J.M., Rohrer, J.D., Alexander, D.C., 2018. Uncovering the heterogeneity and temporal complexity of neurodegenerative diseases with Subtype and Stage Inference. Nat Commun 9, 4273. https://doi.org/10.1038/s41467-018-05892-0

Zhang, T., Koutsouleris, N., Meisenzahl, E., Davatzikos, C., 2015. Heterogeneity of Structural Brain Changes in Subtypes of Schizophrenia Revealed Using Magnetic Resonance Imaging Pattern Analysis. Schizophr Bull 41, 74–84. https://doi.org/10.1093/schbul/sbu136

Zhang, W., Deng, W., Yao, L., Xiao, Y., Li, F., Liu, J., Sweeney, J.A., Lui, S., Gong, Q., 2015. Brain Structural Abnormalities in a Group of Never-Medicated Patients With Long-Term Schizophrenia. Am J Psychiatry 172, 995–1003. https://doi.org/10.1176/appi.ajp.2015.14091108

Zhang, X., Mormino, E.C., Sun, N., Sperling, R.A., Sabuncu, M.R., Yeo, B.T.T., the Alzheimer’s Disease Neuroimaging Initiative, 2016. Bayesian model reveals latent atrophy factors with dissociable cognitive trajectories in Alzheimer’s disease. Proc Natl Acad Sci USA 113, E6535–E6544. https://doi.org/10.1073/pnas.1611073113

Zhirong Yang, Oja, E., 2010. Linear and Nonlinear Projective Nonnegative Matrix Factorization. IEEE Trans. Neural Netw. 21, 734–749. https://doi.org/10.1109/TNN.2010.2041361

Zhu, J., Zhuo, C., Liu, F., Xu, L., Yu, C., 2016. Neural substrates underlying delusions in schizophrenia. Scientific Reports 6, 33857. https://doi.org/10.1038/srep33857

Zhuo, C., Ma, X., Qu, H., Wang, L., Jia, F., Wang, C., 2016. Schizophrenia Patients Demonstrate Both Inter-Voxel Level and Intra-Voxel Level White Matter Alterations. PLOS ONE 11, e0162656. https://doi.org/10.1371/journal.pone.0162656

